# BrainPhys neuronal medium optimized for imaging and optogenetics in vitro

**DOI:** 10.1101/2020.09.02.276535

**Authors:** Michael Zabolocki, Kasandra McCormack, Mark van den Hurk, Bridget Milky, Andrew Shoubridge, Robert Adams, Jenne Tran, Anita Mahadevan-Jansen, Philipp Reineck, Jacob Thomas, Mark R Hutchinson, Carmen Mak, Adam Añonuevo, Leon Harold Chew, Adam J. Hirst, Vivian M. Lee, Erin Knock, Cedric Bardy

## Abstract

The capabilities of imaging technologies, fluorescent sensors, and optogenetics tools for cell biology have improved exponentially in the last ten years. At the same time, advances in cellular reprogramming and organoid engineering have quickly expanded the use of human neuronal models *in vitro*. Altogether this creates an increasing need for tissue culture conditions better adapted to live-cell imaging. Here, we identified multiple caveats of traditional media when used for live imaging and functional assays on neuronal cultures (e.g., phototoxicity, suboptimal fluorescence signals, and unphysiological neuronal activity). To overcome these issues, we developed a new neuromedium, “BrainPhys™ Imaging”, in which we adjusted fluorescent and phototoxic compounds. The new medium is based on the formulation of the original BrainPhys medium, which we designed to better support the neuronal activity of human neurons *in vitro* ^1^. We tested the new imaging-optimized formulation on human neurons cultured in monolayers or organoids, and rat primary neurons. BrainPhys Imaging enhanced fluorescence signals and reduced phototoxicity throughout the entire light spectrum. Importantly, consistent with standard BrainPhys, we showed that the new imaging medium optimally supports the electrical and synaptic activity of midbrain and human cortical neurons in culture. We also benchmarked the capacity of the new medium for functional calcium imaging and optogenetic control of human neurons. Altogether, our study shows that the new BrainPhys Imaging improves the quality of a wide range of fluorescence imaging applications with live neurons *in vitro* while supporting cell viability and neuronal functions.

## Introduction

For many years, light microscopy was restricted to anatomical and cellular morphology analysis. In the last two decades, technological advances in fluorescent probes, optogenetics, camera sensors, light-emitting diodes, and microscopes have unleashed the capacity of light microscopy for functional imaging ^2–8^. Calcium imaging is an increasingly popular electrophysiological assay ^9^. Neuroscientists are slowly replacing laborious patch-clamping experiments with optical voltage sensors ^7, 10, 11^. Precise light-emitting molecular pH sensors provide insights into synaptic or lysosomal mechanisms. At the same time, light has also become a powerful tool to remotely control the function of neurons. Optogenetics allow precise activation and inhibition of subtypes of neurons ^8, 12, 13^. The design of photo-releasable caged molecules advances our understanding of molecular physiology ^14–16^. However, unintended light-induced cytotoxicity with live-cell imaging technologies *in vitro* may hinder scientific discoveries ^17^. Limiting the intensity and duration of light exposure or using longer wavelengths (>500 nm) may partially reduce live imaging-induced cytotoxicity ^18–21^. However, it does not resolve the issue and comes at the cost of sub-optimal signal-to-background ratios.

Neuronal cultures are notoriously difficult to maintain and have a low tolerance for manipulation and stress. The most likely cause of light-induced toxicity is the generation of reactive oxygen species produced by light-reactive components in the media ^22–24^. Previous studies have shown that adjusting phosphate buffers such as N-(2-hydroxyethyl)piperazine-N’-2-ethane sulfonic acid (HEPES) may reduce the accumulation of light-induced cytotoxic products in tissue culture media ^25^; further removing riboflavin from Neurobasal™- and DMEM™-based media improved rat primary neuronal cell survival when exposed to blue light (470 nm)^26^. These base media optimized for phototoxicity may help with some imaging experiments. However, we found that with human neuronal models, DMEM and Neurobasal are suboptimal for the firing of action potentials and synaptic activity (glutamatergic and GABAergic), which is a major caveat to model the physiology of the human brain *in vitro* ^1^.

Thus, there is a need for a new neuronal medium specialized for functional imaging experiments. Such a medium needs to support neurophysiological activity, minimize light-induced toxicity, and enhance fluorescent signal-to-background ratios. To address this need, we implemented the formulation of the original BrainPhys™ (BP) medium ^1^ in the design of a new functional imaging neuromedium, which we called BrainPhys™ Imaging (BPI). We benchmarked its performance in comparison with other media currently available. We found that removing the light-reactive components from BP enhanced fluorescent signals and reduced neuronal phototoxicity. We demonstrated that the new formula could be used to culture neurons without a detrimental effect on cell survival and its osmolality match physiological levels of human CSF. Finally, we used patch-clamping, multi-electrode arrays, calcium imaging, and optogenetics to show that the formulation of BrainPhys™ Imaging supports optimal electrophysiological functions of human neurons *in vitro*.

## RESULTS

### Adjustment of standard BrainPhys basal medium formula to optimize live-cell imaging

To optimize the original BrainPhys basal medium for imaging, we adjusted the components within the formulation which we identified as potentially increasing fluorescent background noise and phototoxicity. The major changes made were the removal of phenol red, and adjustments of the vitamins (e.g., riboflavin) and pH buffers. The vitamin content of the medium appeared as the main factor responsible for phototoxicity and autofluorescence interference at excitation wavelengths less than 500 nm. The adjustments made had no significant effect on the overall osmolality of the medium (**Figure 1A**). This is an essential attribute of BrainPhys that we wished to maintain as it is similar to that of human cerebrospinal fluid (~300 mOsmol/L). We also confirmed that the osmolality of media conditioned with hiPSC derived neurons was identical to the fresh media and stable over time (tested for up to 2 weeks, **Figure S1A**). The pH was also maintained at physiological levels (pH 7.4) in the new formulation. The new formulation is referred to as BrainPhys Imaging (BPI). For comparison, we also evaluated two other commercially available imaging media, BrightCell™ NEUMO and FluoroBrite™ DMEM, which were previously modified based on Neurobasal and DMEM, respectively ^26^. We previously demonstrated that the more physiological composition of BrainPhys basal medium improves the electrophysiology and synaptic activity of human neurons *in vitro* in comparison to DMEM and Neurobasal ^1^. In the present study, we compared the imaging performance, phototoxicity, and electrophysiological properties of cells in BrainPhys Imaging to standard neuronal media and other imaging media currently available.

**Figure 1.**
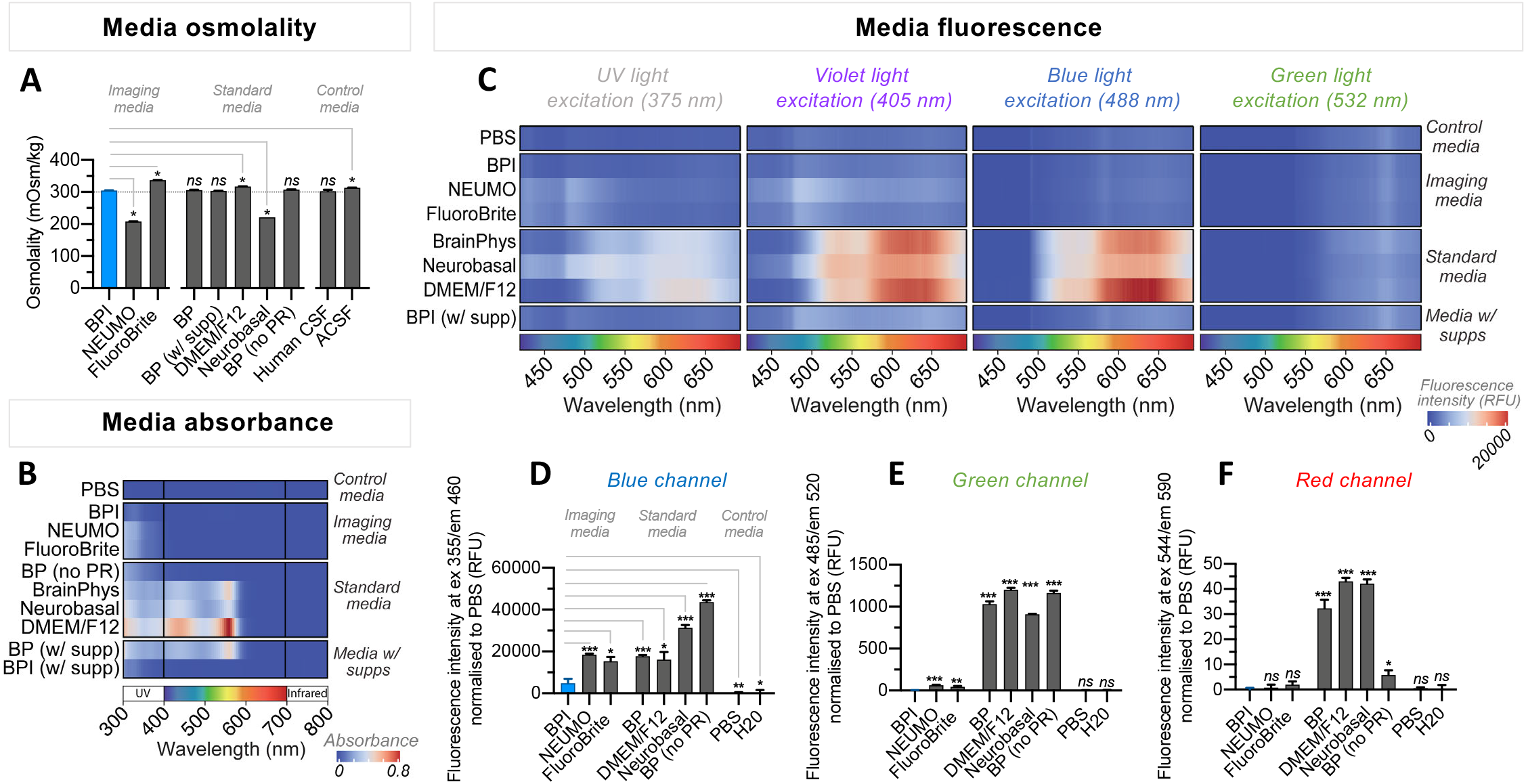
BrainPhys Imaging reduces auto-fluorescence and light absorbance. **A,** shows that the osmolality of BrainPhys Imaging (BPI) is around 305 mOsmol/kg, which is similar to physiological human cerebrospinal fluid (hCSF). Neurobasal and NEUMO were significantly lower (210-220 mOsmol/kg) and DMEM/F12 and FluoroBrite higher (317-338 mOsmol/kg). Data was collected from 3-4 replicates per condition. **B-F,** compares the optical properties of BPI with other basal media specialized for imaging (NEUMO, FluoroBrite), standard basal neuro-media (BrainPhys “BP”, BrainPhys without phenol red “BP (w/out PR)”, DMEM/F12, Neurobasal) and control media (phosphate-buffered saline (PBS) and deionized water (H_2_O)). **B,** shows the absorbance spectra from 300-800 nm acquired from basal media alone (without cells), and after adding supplements required for the long-term culture of brain cells. BPI shows lower absorbance than all other culture media tested. Adding supplements to BPI did not increase the absorbance, which was tested between 300-800 nm. Virtually all fluorophores used for cell imaging require light stimulation above 300 nm. **C-F,** reveals the mean auto-fluorescence intensities of basal media (without cells) across the entire visible spectrum. BPI shows auto-fluorescence intensities similar to PBS. **C,** shows the emission spectra across 400-700 nm captured for the 375 (violet), 405 (blue), 488 (green) and 532 nm (red) excitation wavelengths from test and control media. **D-F,** autofluorescence at 460, 520, and 590 nm emission wavelengths were measured following excitation at 355, 485 and 544 nm, respectively. Results were generated from eight replicate wells per medium across three independent experiments. For normalization, the mean fluorescence intensity in PBS was subtracted from the other media. Values represent mean ± SEM. Significance determined via two-tailed non-parametric unpaired (Mann Whitney) tests. P-values are annotated as follows: * for P<0.05, ** for P<0.01, and ns for P>0.05.

### BrainPhys Imaging has low absorbance and autofluorescence across the visible light spectrum

We compared the performance for fluorescence imaging of BPI to other tissue culture media (**Figure 1, 2**). First, we measured the absorbance spectrum (300 nm – 800 nm) of BPI basal and other basal media without supplements (**Figure 1B**). The absorbance above 600 nm was close to zero for all media tested. However, below 600 nm the relative level of absorbance differed between basal media. All classical basal media used for neuronal culture (standard BrainPhys, Neurobasal, and DMEM/F12) had relatively high absorbance for all wavelengths below 600 nm. Removing phenol red from standard media helped to reduce the absorbance between 400 nm to 600 nm. Further adjustment of vitamins and pH buffer in BPI also helped to reduce the absorbance below 400 nm to levels almost as low as PBS. We also confirmed that the addition to the basal medium of a range of supplements (SM1, N2-A, BDNF, GDNF, Ascorbic Acid, cAMP, Laminin; selection based on the standard BrainPhys formulation established in ^1^) did not increase the absorbance above 300 nm (**Figure 1B**).

**Figure 2.**
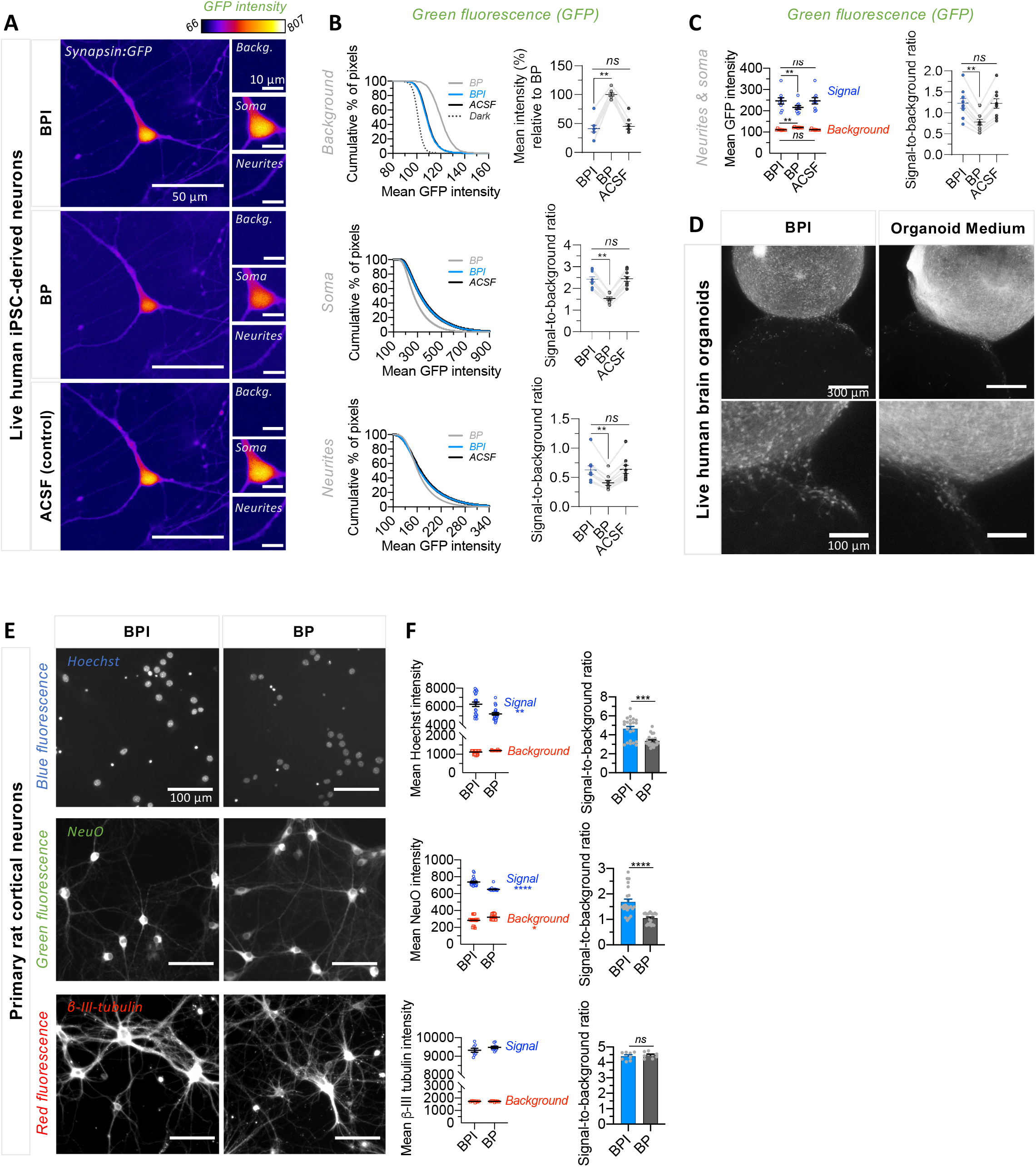
BrainPhys Imaging increases the fluorescence signal-to-background ratio of neuronal cultures. Fluorescent imaging in BPI medium improves signal-to-background ratios relative to BrainPhys medium. For signal-to-background ratio quantification, mean intensity ‘signal’ values at nuclei (Hoechst), combined neurites and soma (NeuO, β-III tubulin, GFP), or isolated neurite and soma (GFP) regions of interest (ROI) were measured and compared to mean ‘background’ intensities. For each field-of-view (FOV), signal-to-background ratios were calculated using the following formula: [(signal intensity – background intensity)/background intensity]. **A-C,** BPI improves signal-to-background ratios at soma and/or neurite regions and reduces background intensities relative to BP. **A,** representative fluorescent images of live human iPSC-derived neurons expressing GFP (excitation/emission peaks: 488/509 nm; filters: 470±40/525±50 nm) and imaged in BPI, BP or artificial cerebrospinal fluid (ACSF) using a 40x water immersion lens (0.8 NA). Higher mean GFP intensities at soma and neurite regions were observed in BPI and ACSF relative to BP, whereas background intensities were reduced. **B-C,** the analysis was conducted on GFP images from 9 field-of-views of the same cells in BrainPhys, BPI or ACSF. All data is paired. **B, left:** shows the cumulative percentage (%) of pixels at background, soma and neurite regions in BP (grey), BPI (blue), and ACSF (black) above a range of mean GFP intensity thresholds. Camera dark counts without medium and cells are labeled as ‘dark’ (top panel). **B, right:** shows that mean intensity values at background regions (top panel) in ACSF and BPI were significantly reduced relative to BP. Signal-to-background ratios at isolated neurite and soma regions were significantly higher in BPI and ACSF than BP. Combining neurite and soma regions in **(C)** also showed improved signal-to-background ratios in BPI and ACSF compared to BP. **D,** representative images of brain assembloids representing a GFP-labelled ventral organoid fused with a non-labeled dorsal organoid imaged using 488nm laser light in Organoid Maturation medium (right) or BPI (left). Note the increased visibility of interneuron migration in the organoids imaged in BPI (bottom). Images in both media are displayed with the following minimum/maximum intensity counts: 0/75. **E,** shows representative fluorescent images of primary cortical neurons stained and imaged live with Hoechst 33342 (excitation/emission peaks: 350/461 nm; filters: 377±25/447±30 nm), live with NeuroFluor NeuO (470/555 nm; 475±17/536±20 nm), and fixed with β-III tubulin-Dylight594 (594/618 nm; 562±20/624±20 nm). Images taken using a 20x air immersion lens (0.45 NA) in both media are displayed with the following minimum/maximum intensity counts: 0/8500 (Hoechst), 0/3600 (NeuO), 1000/12000 (β-III tubulin). **F,** signal-to-background ratios were improved when imaging Hoechst in BPI (n=23 FOVs) compared to BP (n=24 FOVs). This was also seen for NeuO images captured in BPI (n=23 FOVs) relative to BP (n=22 FOVs). No significant differences were found when imaging β-III tubulin in either BPI (n=8 FOVs) or BP (n=9 FOVs). Images were collected from two biologically independent experiments (Hoescht/NeuO; β-III tubulin) with one well per condition. See also **S1.** All values represent mean±SEM. **B, C,** significance determined via two-tailed non-parametric paired (Wilcoxon) tests and **(F)** non-parametric unpaired (Mann Whitney) tests. P-values are annotated as follows: * for P<0.05, ** for P<0.01, **** for P<0.0001 and ns for P>0.05.

Then, we measured the relative autofluorescence of each media in the absence of cells (**Figure 1C, D - F**). We used a custom-made bifurcated fiber probe that we positioned within the media. We then excited each medium at four wavelengths sequentially from violet to green (375, 405, 488, 532 nm) and measured the emission spectra from 400 to 700 nm. We found that BPI basal medium autofluorescence across the visible light spectrum was as low as for PBS (**Figure 1C**). In contrast, all other standard media tested (BP, DMEM, Neurobasal) showed much higher autofluorescence signals at excitation wavelengths from violet to green. When tested with short wavelengths light excitation (375 nm, 405 nm), BPI also showed slightly lower autofluorescence than NEUMO and FluoroBrite (**Figure 1C**). Low autofluorescence of BPI across the light spectrum was maintained despite adding supplements. For further statistical comparison, we repeated these experiments using a plate reader **(Figure 1D - F)**. We compared the mean fluorescence intensity with three dichroic filters: blue (excitation/emission at 355/460 nm), green (excitation/emission at 485/520 nm), and red (excitation/emission at 544/590 nm). The blue, green, and red autofluorescence levels of BPI were significantly lower than that of any other standard basal neuromedia (BrainPhys, DMEM, Neurobasal) **(Figure 1D - F)**. At shorter wavelengths (blue and green), BPI also outperformed the autofluorescence of other specialized imaging media (BrightCell™ NEUMO and FluoroBrite™ DMEM) **(Figure 1D, E)**. At longer wavelengths (red), the autofluorescence of BPI and other imaging media was similar **(Figure 1F)**. The green and red autofluorescence of BPI was equivalent to that of PBS and H_2_O **(Figure 1E, F)**. However, we noted that the blue autofluorescence in BPI remained slightly higher than PBS or H_2_O, despite outperforming all the other media **(Figure 1D)**. The removal of phenol red alone did not show any reduction in autofluorescence in the blue and green channels but helped to lower the red autofluorescence **(Figure 1D-F)**. Autofluorescence with light excitation at a longer wavelength (532 nm) was relatively low for all media tested (**Figure 1C**), though the plate reader experiments revealed a relatively small but significant reduction in autofluorescence for BPI compared to standard media in the red channel (**Figure 1F**). Overall, the absorbance and autofluorescence of BPI were most distinctively reduced for excitation wavelengths between ultraviolet and blue compared to other neuronal media (**Figure 1B, C**). Most importantly, the modifications made in BPI lowered the absorbance and autofluorescence at levels similar to PBS across all visible wavelengths.

### BrainPhys Imaging improves fluorescent signal-to-background ratios

We then showed that the lower level of autofluorescence of BPI enhanced signal-to-background ratios when imaging brain cells *in vitro*, using healthy human iPSC-derived neurons in monolayers (**Figure 2A-C, S2, S3**) and organoids (**Figure 2D**) or rat primary cortical neurons (**Figure 2E, F, S1C-F**). Live human iPSC-derived neurons were infected with lentivectors to express green (eGFP) using a synapsin promoter (**Figure 2A-C, S2 and S3**). The human neurons were imaged live in standard BrainPhys or BPI with identical imaging parameters. BrainPhys Imaging significantly increased signal-to-background ratios at the neurite and soma regions, and significantly reduced mean background intensities (**Figure 2A-C**). When combining analysis at soma and neurite regions, similar increases in signal-to-background ratios were observed. Re-perfusing the cells with ACSF, after testing them subsequently in BPI and BP, recovered signal-to-background ratios in the background, soma and/or neurite regions to levels identical to BPI (**Figure 2A-C**). This confirmed that the effect was not simply due to possible photobleaching. Overall, all the neuronal monolayer and organoid images in BPI showed increased signal-to-background ratios for lower wavelengths (blue and green) compared to all other media tested (standard BrainPhys, **Figure 2A-C, E, F**, **and Figure S1C-F, S2, S3;** Organoid medium, **Figure 2D;** BrainPhys Without Phenol Red and NEUMO, **Figure S1C-F**). The live imaging of an assembly of brain-region specific organoids (assembloids, ^27^) in BPI also allowed for better visualization of migrating neurons than in standard organoid maturation medium (**Figure 2D**). We then used rat cortical primary neurons to test the signal-to-background ratio at different wavelengths. The primary neurons were either fixed and labeled in red with β-III-tubulin and DyLight™ 594 (excitation/emission at 593/618 nm) or live in green with the neuron-specific dye NeuroFluor™ NeuO (excitation/emission at 468/557 nm). For Hoechst™ 33342 (excitation/emission at 361/497 nm), the neurons were either labeled live or fixed first. For each fluorophore, all images were conducted with identical imaging parameters but in different media (**Figure 2E, F and S1C-F)**. For longer wavelengths (red), we found no significant improvement in signal-to-background ratio with images taken in BPI (**Figure 2E, F and Figure S1E, F**).

Overall our data supports that BPI constitutes a new alternative imaging neuronal medium that provides lower absorbance, lower autofluorescence, and enhanced fluorescent signal-to-background ratios throughout the entire light spectrum from ultraviolet (300 nm) to infrared (800 nm), which is used by virtually all fluorophores, sensors and imaging probes in cell biology.

### BrainPhys Imaging reduces phototoxicity across the light spectrum (from violet to red)

To compare the extent of phototoxicity on neuronal cultures in different media, we first measured the viability of primary rat cortical neurons after prolonged exposure to violet (395-405 nm; 430 Lux ± 3.70 S.E.M), blue (450-475 nm; 14 Lux ± 1.41 S.E.M) and red (620-740 nm; 55 Lux ± 4.99 S.E.M) LED light inside a temperature-controlled incubator **(Figure 3A-D)**. Neurons exposed to light in phototoxic media exhibited broken neurites and disintegrated cell bodies (**Figure 3A-C; Figure S5);** the number of live cells was counted post light-exposure to quantify cell viability. Exposure to violet and blue light caused cell death in standard BrainPhys medium (with or without phenol red) but not in BPI **(Figure 3A-D).** We also compared the phototoxicity of the three wavelengths in BPI to NEUMO. Although NEUMO basal is provided with a supplement (SOS), we were unable to sustain healthy neurons with this combination for long enough to perform this assay **(Figure S4B).** Thus, we tested NEUMO basal medium combined with the same supplement that was added to BrainPhys basal in these experiments (SM1 Neuronal Supplement). In these conditions, NEUMO performed comparably to BPI when stimulated for 12h with blue light **(Figure 3D).** However, when using higher energy violet light (430 Lux ± 3.70 S.E.M) for 6h, the cells only survived in BPI **(Figure 3D)**. Red light did not appear phototoxic in any media we tested, even when exposed for longer at four times the Lux intensity of the blue light test **(Figure 3D)**. To quantify more subtle phototoxic effect preceding cell death, we exposed BP and BPI media to ambient light in the tissue culture hood for 24h and fed human PSC-derived midbrain and cortical neurons daily for up to 7 days (half media change) while measuring the accumulation of lactate dehydrogenase (LDH) in the supernatant. LDH is a cytosolic enzyme, which is present in neurons and released into the supernatant upon damage to the plasma membrane. We found that media exposure to ambient light triggered a higher release of LDH in cultures fed with BP compared to BPI within two days (**Figure 3E, F**). These results were consistent both with cortical and midbrain human neurons (**Figure 3E, F**). Riboflavin, which is present in BP and other classical neuromedia, can release free radicals such as peroxide hydrogen (H_2_O_2_). We found that H_2_O_2_ levels significantly increased in BP after 24h exposure to ambient light but not in BPI. This H_2_O_2_ increase was observed despite adding antioxidants in supplements to both media (**Figure 3G**). The spectrum of the ambient light from our tissue culture hood showed multiple-wavelength peaks between 400 nm and 650 nm (**Figure 3H**). Therefore, we used an LED controller instead of ambient light to show that wavelengths around 475 nm (**Figure 3I**, **S4A**) were sufficient to increase the level of H_2_O_2_ in BP medium in a dose-dependent manner (**Figure 3J**). At the highest dose tested (~110mW at 475 nm) the level of H_2_O_2_ doubled in BP and only slightly increased in BPI. Oxidative stress is also often associated with mitochondrial impairments. Indeed, we observed a significant reduction of the mitochondrial potential of neurons exposed to violet light in standard BP (**Figure 3K**). This effect appeared to precede neuronal death (**Figure S4C**) and was comparable to exposing the neurons to 100uM of H_2_O_2_. In contrast, the mitochondrial potential of neurons was not affected by light stimulation in BPI (**Figure 3K**). Altogether, BPI was the only neuronal medium tested that showed no neuronal phototoxicity across the full visible light spectrum.

**Figure 3.**
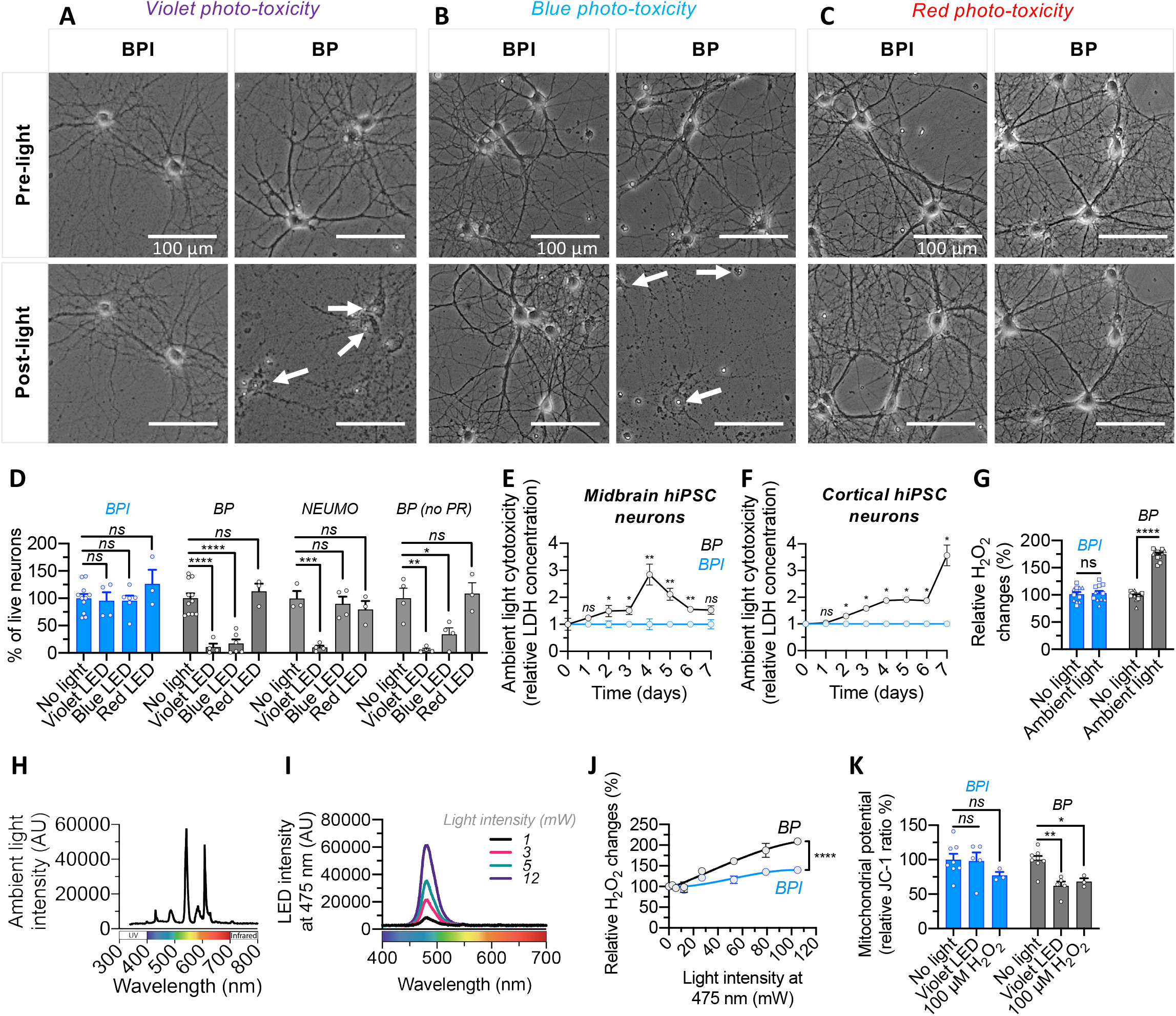
BrainPhys Imaging reduces photo-toxicity. Phototoxicity was reduced in BPI compared to standard BP, BP without phenol red (BP no PR), and NEUMO. **A-C,** representative images of rat cortical primary neurons cultured in standard BP and BPI before and after exposure to violet, blue, and red LED light. See also *figure S5*. Cell exposure to a violet LED was completed over 6 hours, blue LED over 12 hours and a red LED over 18 hours. Only neurons cultured in BPI maintained healthy morphology post-light exposure for violet and blue LEDs. White arrows highlight dying neurons post-light exposure. Red LED had no apparent phototoxicity. **D,** shows the percentage loss of primary rat cortical neurons following violet, blue and red LED exposures in BP, BPI, NEUMO and BP (no PR) media. Data were collected across 3-5 independent experiments, each including 75 field-of-views analyzed across three replicate wells. The number of neurons (per cm^2^) post-light exposure were normalized to the number of neurons (per cm^2^) in ‘no light’ conditions. **E-F,** BPI and BP media were placed under ambient tissue culture hood light for 24h before feeding the cells. When exposed to ambient light, BP induced significantly more cytotoxicity in human midbrain and cortical neurons than BPI. The same midbrain or cortical supplements were added to both BP and BPI basal media. Cytotoxicity was measured with relative changes in LDH over 7 days, (quantified in supernatant collected from neurons with the absorbance of Formazan at 490 nm). Data was collected from 3-6 replicate wells per day. **G,** reactive oxygen species (H_2_O_2_) in BP was significantly increased following 24-hour stimulation with ambient light but not in BPI. Per condition, data was collected from 12 replicate wells – 6 with cortical supplements (triangles) and 6 with midbrain supplements (squares). **H,** ambient light emission spectrum in tissue culture hood used for experiments. **I,** blue LED light emission spectrum used in ‘J’. **J,** relative H_2_O_2_ levels of BPI and BP basal media (without cells) after 24-hour stimulation with 10 × 5 ms flashes of blue LED light (475nm) at 10 Hz and a 30 second inter-burst interval. Incremental blue LED power intensities were used. For each condition and intensity, data was collected from a total of 6 replicate wells. H_2_O_2_ levels were normalized to ‘no light’ conditions. See *also Figure S4A and C.* **K,** primary rat cortical neurons in BPI and BP media supplemented with SM1 were exposed to violet LED light inside a temperature-controlled incubator (37°C) for 1 hour or treated with 100 μM of H_2_O_2_. Mitochondrial health was assessed with a mitochondrial membrane potential indicator (JC-1). In BP, mitochondrial health was significantly reduced following violet LED light exposure or H_2_O_2_ treatment. Only neurons exposed to BPI maintained mitochondrial health. Data were collected across 3-5 independent experiments, across 3 biological replicates. Results are shown normalized to ‘no light’ conditions. *See also S4C*. Values represent mean ± SEM. **D,** Significance determined via unpaired T-test, (**E-G,K**) two-tailed non-parametric unpaired (Mann Whitney) tests and **(J)** sum-of-squares F-test. P-values are annotated as follows: * for P<0.05, ** for P<0.01, *** for P<0.001, **** for P<0.0001 and ns for P>0.05.

### BrainPhys Imaging supports optimal action potentials and synaptic activity in human neurons

We then asked whether BPI maintains the fundamental electrophysiological properties of human neurons. We used patch-clamping and MEA to test the effect of BPI on the electrophysiological activity of human neurons (**Figure 4**). The neurons were matured from human PSC-derived neural progenitors in standard BrainPhys for 4 to 5 months before patch-clamping (**Figure S6A**). The cells were selected for patch-clamping based on their neuronal morphology visualized with synapsin:GFP lentiviral marker (**Figure S6B**). During the whole-cell recordings, we alternated the perfusion between BPI and artificial cerebrospinal fluid (ACSF), which is used as the gold-standard for acute electrophysiology. We first found that acute perfusion of BPI had no negative effect on the types of action potential (AP) firing patterns, which were defined previously in ^28^ (AP Types 1-5; **Figure S6C**). To confirm that the new medium does not have more subtle effects, we further examined the electrophysiological properties of “mature” neurons (Type 5). Our results show that the cellular mechanisms critical to generate action potentials were healthy in BPI and we found no difference in the electrophysiology of the cells when patched in ACSF (**Figures 4, S6**) or in standard BP (**Figure S7**). To measure evoked action potentials (in current-clamp) or voltage-gated sodium and potassium currents (in voltage-clamp), we maintained the resting potential at −70 mV (by injecting on average 50 pA of current) and applied incremental depolarizing steps of current/voltage (**Figure 4A, Figure S6D**). Voltage-gated sodium/potassium currents, rheobase, AP amplitude, AP frequencies, and AP hyperpolarizing amplitudes were similar when the recording was made in BPI or in ACSF (**Figure 4C-I**). When no current was injected in the patch-clamped neurons, their resting membrane potential was around −50 mV and their AP threshold approached physiological levels (−40mV) both in BPI and ACSF **(Figure 4C)**. No significant differences were found between the membrane resistance and capacitance of neurons patch-clamped in either BPI or ACSF (**Figure 4J, K**). Our results demonstrate the capacity of human neurons to generate optimal action potentials in BPI.

**Fig 4.**
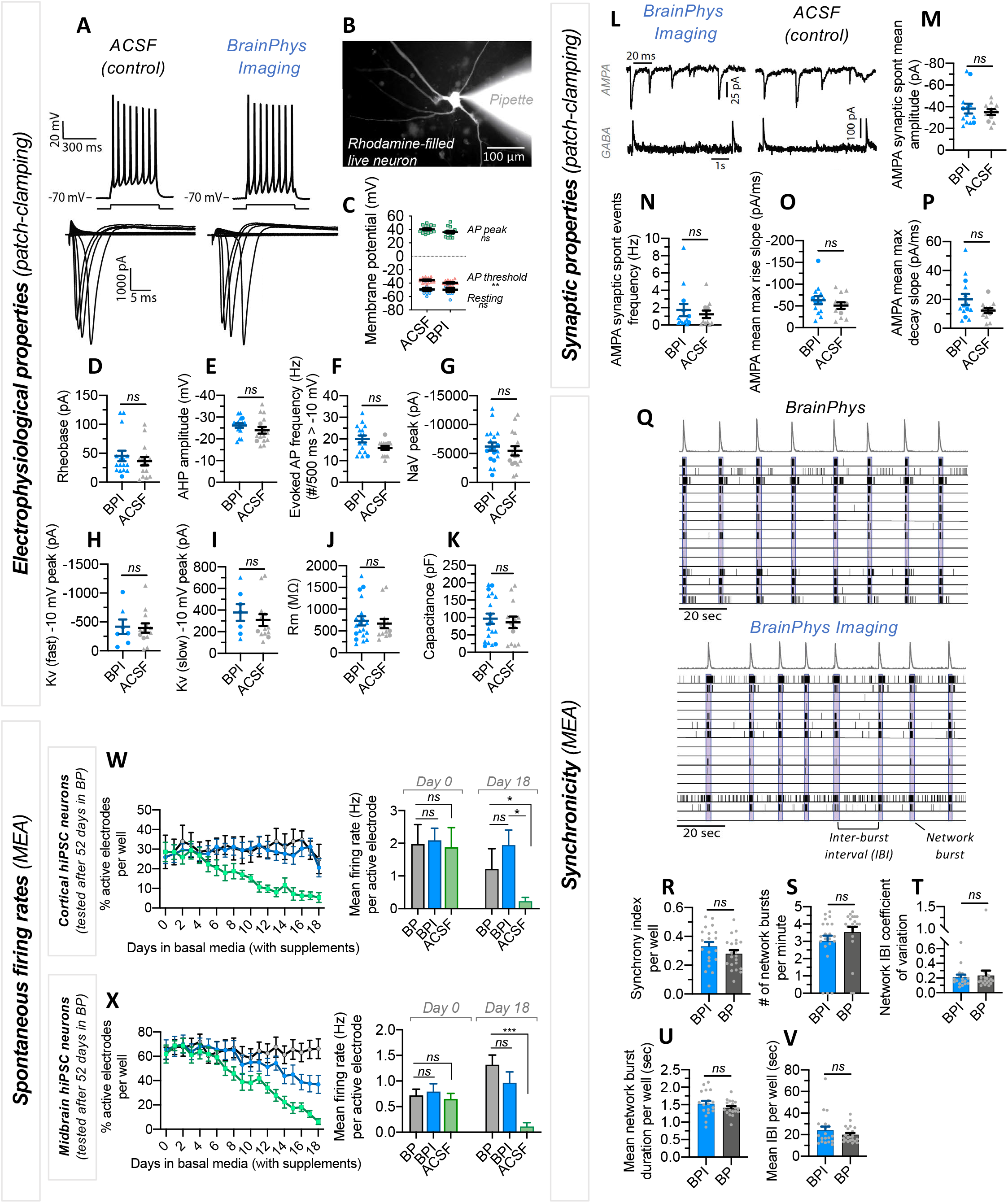
BrainPhys Imaging supports optimal electrophysiological properties of human cortical and midbrain neurons *in vitro*. **A-P**, single-cell patch-clamp recordings of human pluripotent stem cell-derived neurons matured in standard BP + supplements medium for >12 weeks, and patch clamped in BPI or ACSF media. All patch clamped cells (n=35) included in the analysis were classified as Type-5 neurons (evoked APs > 10 Hz, amplitudes > −10 mV) based on methods previously established in Bardy et al. 2016. See also Figure S6. A subset of neurons (n=6) was recorded in both media, but the majority were recorded in only one medium, therefore, unpaired statistics were performed. All neurons were patched from a total of 18 coverslips. Each point on the graphs represents a single neuron. **A, top:** typical evoked action potential (AP) traces following a 500 ms depolarizing current step when patched in ACSF or BPI. **Bottom:** current-voltage characteristics (I-V curve) with +5 mV current steps increments from rest at −70 mV. Voltage-dependent sodium (Nav) and potassium (Kv) current traces shown, respectively, below and above the x-axes. **B,** shows a representative image of a single neuron filled with rhodamine following patch-clamp recordings. **C-E,** action potential properties were similar in BPI (n=15 neurons) and ACSF (n=15 neurons). **C,** pooled data summarizing peak AP amplitudes, threshold and resting membrane potential values. Resting membrane potential was measured when 0 pA current was injected. **D,** rheobase represents the minimum depolarising current step required to evoke an AP. **E,** peak afterhyperpolarization (AHP) amplitudes following an AP. **F,** the maximum firing frequency of AP evoked by 500ms depolarization steps (only spikes with amplitudes > −10 mV were included to measure the firing frequency). **G-I,** current-voltage characteristics were similar in BPI (n=19 neurons) and ACSF (n=16 neurons). **G,** peak NaV current amplitudes. **H-I,** peak amplitudes of rapidly and slowly inactivating Kv currents. **J-K,** membrane resistance (Rm) and capacitance values were similar in BPI (n=19 neurons) and ACSF (n=13 neurons). **L-P,** synaptic events mediated by AMPA receptors (excitatory postsynaptic currents, ePSCs) and GABAa receptors (inhibitory postsynaptic currents, iPSCs) were similar in BPI (n=14 neurons) and ACSF (n=12 neurons). **L,** typical spontaneous ePSC (top) and iPSC traces (bottom). **M-P,** average properties of spontaneous synaptic events recorded for 4 min. **M,** mean ePSC amplitudes. **N,** mean ePSC event frequency. **O, P,** max rise and max decay slopes of ePSCs. Symbols represent human neurons tested first (triangles) or second (circle) in either medium **(D-K, M-P)**. **Q-V,** midbrain human iPSC-derived neurons were matured for 100 days in BP before switching to BP or BPI for 2 hours. The same midbrain supplements were added to both basal media. Network activity was compared on multielectrode arrays (MEA) across 21 wells per condition (16 electrodes per well), over 7-minute time intervals. **Q,** example 2-minute raster plots of neuronal network activity in BPI or BP show similar network activity (blue boxes) and spike histograms (grey, top). (R-V) Quantification of MEA network activity showed no significant differences between BPI and BP conditions. **W-X,** cortical **(W)** and midbrain **(X)** human iPSC-derived neurons were matured for 52 days in standard BP. Then, the spontaneous action potentials firing rates were compared on multielectrode arrays (MEA) over 18 days in standard BP (black trace), BPI (blue trace) and ACSF (green trace). The same cortical or midbrain supplements were added to all three basal media. Spontaneous MEA activity was recorded for 10 minutes every 24 hours. The activity was calculated from 8-10 wells per condition (16 electrodes per well). The spontaneous firing activity was similar in BP, BPI, and ACSF at first but progressively decreased in ACSF after a few days. In contrast, despite some variability, the activity remained similar in BP and BPI over 18 days. In all graphs, values are shown as mean ± SEM. Significance determined via two-tailed non-parametric unpaired (Mann Whitney) tests. P-values are annotated as follows: * for P<0.05, *** for P<0.001, and ns for P>0.05.

Neural network communication is achieved via synaptic integration and a regulated balance of inhibition and excitation. In the human brain, the neurotransmitter glutamate is responsible for the majority of synaptic excitation, while GABA is responsible for the majority of synaptic inhibition. The summation of excitatory and inhibitory neurotransmitters governs the probability of postsynaptic action potential firing. Hence, we assessed whether BPI supports basic synaptic activity in human neurons. Voltage-clamp recordings from mature human neurons highlighted spontaneous synaptic events mediated by both AMPA and GABA receptors when patched in BPI medium or ACSF (**Figure 4L**). Voltage clamping at the reversal potential of anions (−70 mV) and cations (0 mV) was used to distinguish between glutamatergic (Na^+^ current-mediated) and GABAergic (Cl^−^ current-mediated) events. We found no significant differences in any of the properties of AMPA-mediated synaptic events (amplitude, frequency, kinetics) when mature human neurons were patch-clamped in BPI or in ACSF (**Figure 4M-P**). The functional quality of the neuronal network can also be assessed by measuring the synchronicity of action potential firing on multi-electrode arrays. We found similar synchronicity properties when comparing the activity of human neurons side-by-side either in BPI or BP with the same supplements **(Figure 4Q-V)**. Altogether, our results demonstrate the capacity of human neurons to generate and integrate optimal synaptic communication in BPI.

The electrophysiology of the cells in standard BrainPhys or BPI media is equivalent to gold standard ACSF. However, maintaining neurons in ACSF for more than a few hours can induce signs of neurodegeneration. We also previously showed that ACSF is not adapted to culture neurons for extended periods of time even when adding supplements ^1^. Here we repeated these results by comparing the function of human cortical and midbrain neurons on MEA over 18 days either in standard BP basal, BPI basal or ACSF (**Figure 4W, X)**. In these experiments, we added to all three basal media the same sets of supplements (N2-A, BDNF, GDNF, Vitamin C, cAMP, Laminin, and either SM1 for midbrain or IGF-1 and SM1 without Vitamin A for cortical). The spontaneous electrical activity was similar at first, but after a few days, we observed a significant decline when cultured in ACSF. In contrast, over the same period, the firing activity of cortical and midbrain neurons remained strong in standard BrainPhys medium and BPI (**Figure 4W, X**).

### Functional live-cell imaging of human neuronal culture in BrainPhys Imaging medium

Specific ions such as calcium (Ca^2+^) play essential roles in electrophysiological activity and vesicular release at synaptic terminals and are commonly used as a proxy to measure electrophysiological activity with functional imaging. The level of Ca^2+^ in BP and BPI matches human brain physiological levels (1.1 mM CaCl_2_). We tested the potential of BPI to image the spontaneous calcium activity of live human neurons. We found that functional imaging can be performed in BPI with optimal performance equivalent to neurophysiological salt solution ACSF, and improved performance compared to NEUMO and FluoroBrite imaging media **(Figure 5)**. For signal quantification, we monitored changes in intracellular Ca^2+^ with a calcium-sensitive dye (Fluo-4 AM) while perfusing different media. 1112 regions of interest (ROIs) were selected at the soma of neurons across a total of thirteen field-of-views (FOVs) **(Figure 5A).** The FOVs were imaged over 4 mins at 5 Hz in each of the following perfusates: ACSF, BPI, FluoroBrite, and NEUMO media. To investigate the quality and kinetics of spontaneous calcium activity, we categorized the cells based on the type of detected events into (i) cells with calcium spikes (fast-rising phase events), (ii) cells with calcium waves (slow rising phases), (iii) cells with calcium waves & spikes (combined), or (iv) cells without any spontaneous activity **(Figure S8A)**. BPI did not affect the frequency and amplitude of unitary events (fast-rising phase events, dF/F >5%) in comparison to ACSF **(Figure 5B-D and Figure S8)**. A similar proportion of fast-rising cell types was measured when functional imaging was performed in ACSF compared to BPI **(Figure 5E, F and S8)**. However, a significant decline in the number, proportion, and frequency of fast-rising cell types was observed when the perfusates were switched to NEUMO or FluoroBrite media **(Figure 5I, J, and S9).** For slow rising calcium events, no significant changes in kinetics and cell event types were identified when functional imaging was conducted in BPI compared to ACSF, NEUMO, or FluoroBrite **(Figure 5G, H, K, L, and Figure S8, S9)**. The activity of the majority of these cell types was silenced with a voltage-gated sodium channel antagonist (tetrodotoxin, TTX; 1 μM), demonstrating that the calcium events observed are largely mediated by spontaneous action potential activity **(Figure 5E-G, I, K and Figure S8, S9B-E).** We also alternated randomly the sequence of media tested to eliminate the possibility of order bias. Altogether, our results indicate that BPI provides optimal conditions for functional imaging of live human neurons *in vitro*.

**Fig 5.**
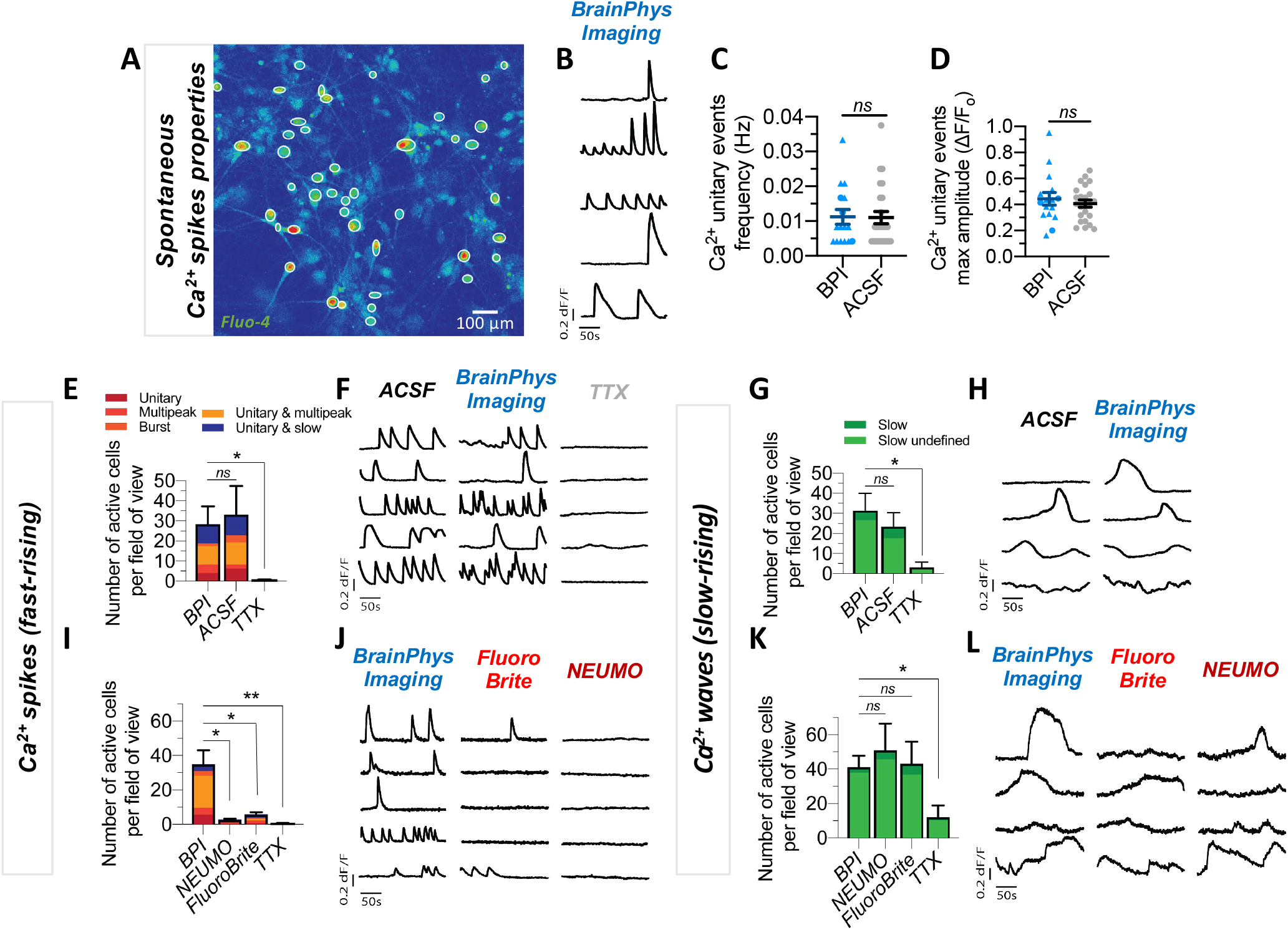
BrainPhys Imaging optimally supports fast-rising calcium spikes in human neurons, *in vitro*. Intracellular Ca^2+^ changes in human pluripotent stem cell-derived neurons were measured with time-lapse image sequences (filmed at 5Hz, 1200 frames over 4 min) of a Ca^2+^ sensor (Fluo-4 AM) under 10x magnification. Time-lapse image sequences were recorded from a total of 1112 human neurons across 13 fields of view (FOVs) from four coverslips. Regions of interest (ROIs) were drawn on the cell soma to determine changes in fluorescence intensity, which were measured as (ΔF/F0), over time. Cells were classified as ‘active’ if at least one event (ΔF/F0 > 5% from baseline) was detected within 4 minutes. Each detected event was manually categorized into fast-rising Ca^2+^ spikes or slow-rising Ca^2+^ waves (see **Figure S8A**). The same FOVs were imaged in multiple perfusates, which included ACSF, BPI, FluoroBrite, and NEUMO. **A,** shows an example of a Fluo-4 fluorescence image taken of a neuronal population in BPI. White circles represent active ROIs. Fluorescent intensity represents intracellular Ca^2+^ levels. **B,** shows typical fast-rising Ca^2+^ spikes from five spontaneously active cells in BPI. **C-D,** shows no significant difference between the average properties of fast-rising unitary Ca^2+^ spike events in BPI (n=16 cells) and ACSF (n=25 cells) from six FOVs across two coverslips. Symbols represent cells recorded first (triangles) or second (circles) in either medium. **E, G,** shows no significant difference between the proportions of cells with fast Ca^2+^ spikes or slow Ca^2+^ waves in BPI and ACSF. The total active cells in BPI (n=238) and ACSF (n=225 cells) were compared across six FOVs from two coverslips. Voltage-gated sodium channel blocker (TTX, 1 μM) abolished virtually all Ca^2+^ spikes and significantly reduced Ca^2+^ waves. **I, K,** comparison of Ca^2+^ signals in BPI, NEUMO, and FluoroBrite shows a significant difference in the proportions of cells with fast Ca^2+^ spikes but no difference in slow Ca^2+^ waves. The cells with Ca^2+^ spikes and Ca^2+^ spikes/waves in BPI (n=243 cells), FluoroBrite (n=39 cells) and NEUMO (n= 18 cells) and the cells with Ca^2+^ waves in BPI (n=288 cells), FluoroBrite (n=302 cells) and NEUMO (n=356 cells) were compared across seven FOVs from two coverslips. **F, H, J, L,** show example traces from the same ROIs in different media. Values are shown as mean ± SEM. Significance determined via two-tailed non-parametric unpaired (Mann Whitney) tests. P-values are annotated as follows: * for P<0.05, ** for P<0.01, and ns for P>0.05.

### BrainPhys Imaging for optogenetics of human neurons

A panoply of optogenetic tools enables remote control of neuronal function with light ^3,8^. To test that BPI is suitable for optogenetics experiments, we transfected human neurons *in vitro* with a lentiviral vector, to drive the expression of channel-rhodopsin tagged with a yellow fluorescent protein (ChETA-eYFP, which is based on ChR2 with two amino acid modifications to improve the kinetics and amplitude of light-evoked currents ^12^). ChETA-eYFP was placed after a human synapsin promoter region (hSyn) to target the expression into neuronal cells. We patch-clamped human neurons expressing ChETA-eYFP (**Figure 6A**). Flashes of blue light (5 ms, 100 ms intervals, at 475 nm) evoked action potentials in BPI comparable to ACSF in all neurons (**Figure 6B**). Using the same light intensity and duration, we found that ten flashes of light at 10 Hz evoked action potentials at a success rate of 99-100% both in ACSF and BPI (**Figure 6C**). The light-evoked conductance of ChETA was similar in BPI and ACSF (**Figure 6C**). Interestingly, when human neurons expressing ChETA-eYFP were patch-clamped in NEUMO and stimulated using the same light parameters (5 ms, 100 ms intervals, at 475 nm), a significant reduction in action potential success rates was witnessed (**Figure 6D and Figure S10**). Furthermore, the action potential light-evoked success rate, peak amplitudes, and the conductance of ChETA remained significantly lower in NEUMO compared to BPI despite increasing LED power (at 475 nm) (**Figure 6E**).

**Fig 6.**
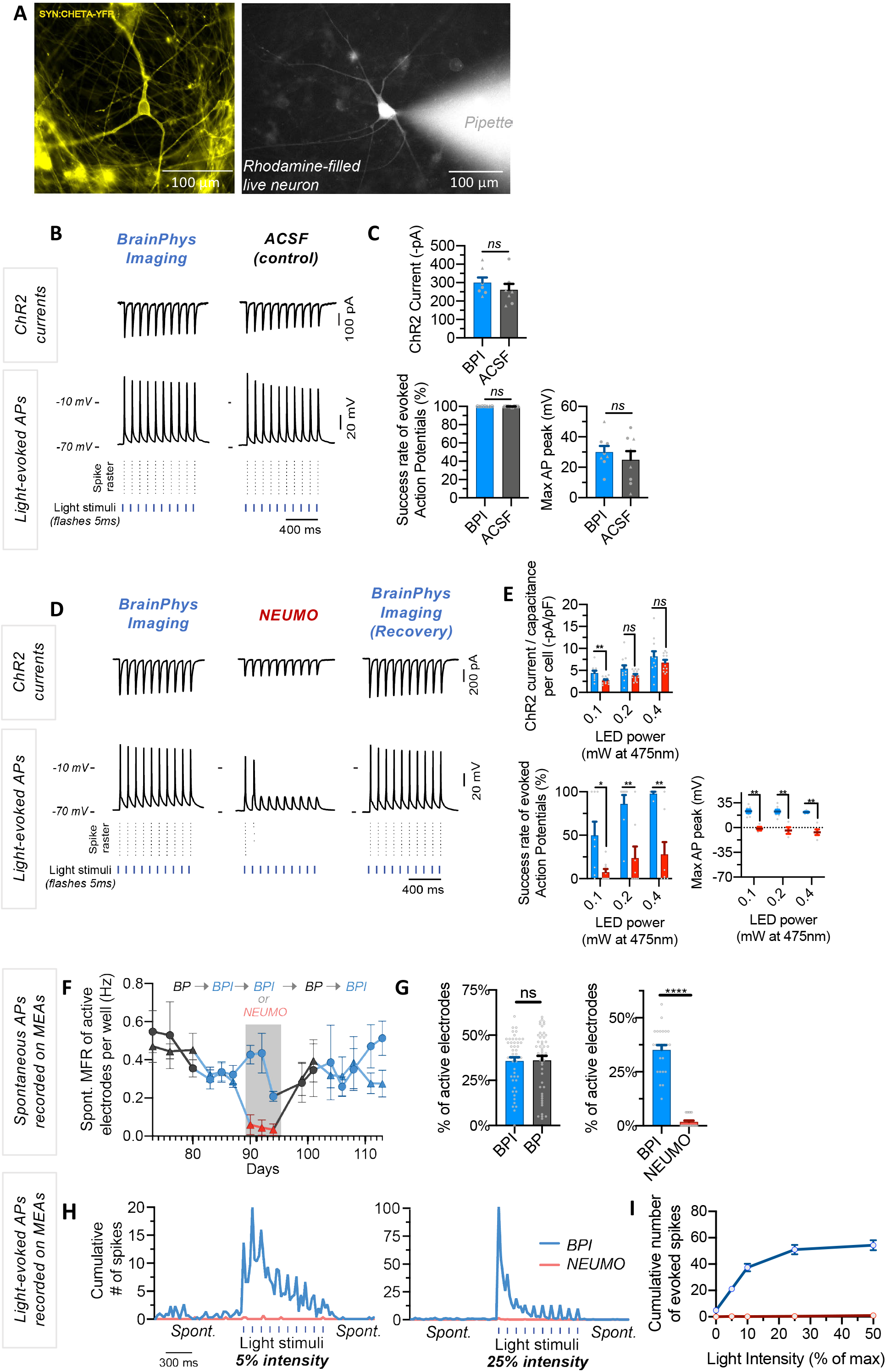
BrainPhys Imaging supports the optogenetic control of human neurons *in-vitro*. **A-E,** optogenetic testing of human iPSC-derived neurons with whole-cell patch-clamp recordings. **A,** live image of typical neurons expressing the optogene synapsin:ChETA-YFP and filled with rhodamine from the patch pipette. **B,** optogenetic responses of patch-clamped human neurons to blue LED stimuli. Whole-cell patch-clamped traces of example neurons stimulated with 0.4 mW 10 × 5 ms flashes of blue light at 10 Hz in BPI and ACSF. Top traces show ChR2-mediated currents recorded in voltage-clamp at −70 mV. Bottom traces show action potential evoked by ChR2-evoked membrane depolarization in current-clamp. Corresponding raster plots highlight consistent light-evoked firing over 10 sweeps. **C,** quantitative comparison shown in the graphs on the right were performed from a total of 8 neurons recorded in ACSF and BPI and tested with identical light stimulation parameters as shown in the trace-examples. Symbols represent human neurons tested first (triangles) or second (circle) in either medium. **D**, optogenetic responses from the same patch-clamped neuron under 0.1 mW of blue light while alternating perfusate from BPI to NEUMO and back to BPI for recovery. **E,** quantitative comparison shown on the graphs on the right were performed from a total of 22 neurons across four coverslips. The perfusate was alternated between BPI and NEUMO. The neurons were stimulated with 10 × 5 ms flashes of blue light at 10 Hz under three different blue LED power intensities (0.1, 0.2, 0.4 mW at 475 nm). **F-I,** optogenetically evoked and spontaneous firing rates of human iPSC-derived neurons expressing synapsin:ChETA-YFP were recorded in a 48-well multielectrode array (MEAs) plate at 37 °C with 5% CO_2_ in either BP, BPI or NEUMO basal media with identical supplements. Spontaneous and light-evoked firing rates were recorded for 10 minutes after feeding. Neurons in this dataset were cultured in standard BP for 82 days before testing different media. 240 electrodes across 15 wells were recorded over 40 days during media testing. **F-G,** recordings collected from one 48-well MEA plate were split into two groups of wells represented in the graph by ‘circles’ (8 wells) and ‘triangles’ (7 wells); Both groups were maintained in BP from neural maturation until they were changed to BPI on day 0. Days 7-13 were the ‘test’ period in which half the wells (7) were switched to NEUMO. From day 14, both groups were cultured in BP to recover for one week and then were switched back to BPI. **H-I,** optogenetics responses of human iPSC-derived neurons recorded with MEAs. The wells in NEUMO and BPI were optically stimulated with blue light (Lumos) at increasing intensities while recording on MEA. Stimulation lasted 1.1s and consisted of 10 flashes of light, each lasting 10ms at 10 Hz. The cumulative number of spikes was summed in binning windows (**H**): 27.5 ms bins; (**I**): 2s bins starting at the onset of the first light stimulus) over three replicates. In (**H)** the cumulative number spikes were plotted with quadratic non-linear curves at various LED intensity (% of max power: 3.9mW/mm^2^). Values are shown as mean ± SEM. Significance determined via two-tailed non-parametric unpaired (Mann Whitney) tests. P-values are annotated as follows: ns for P>0.05.

Our patch-clamping experiments were limited to acute changes of basal media. To test the longer-term effects of the media with supplements, we repeated these experiments with iPSC-derived neurons on a multi-electrode array recorder (MEA). We first recorded the spontaneous electrical activity of neurons in alternative media for one week (three media changes). Overall, despite occasional variability in the level of spontaneous firing, we found no significant difference in the spontaneous activity of the neurons in standard BrainPhys or BPI (**Figure 6F**). In contrast, when neurons were switched to NEUMO for a week, the spontaneous activity dropped and did not recover over time unless it was switched back to BrainPhys medium (**Figure 6F, G**). The human neurons were also transfected with an optogene (LV hSyn:ChETA-EYFP) and stimulated with blue light (475 nm) while on the MEA recorder. Brief flashes of light were able to evoke action potentials in BPI but not in NEUMO (**Figure 6H, I**). Altogether, this data demonstrates that BPI is optimal for optogenetic control of human neurons *in vitro*.

### BrainPhys Imaging supports the viability and function of neuronal cultures

Several components in the standard BrainPhys formulation were reduced or replaced to create BPI. Therefore, we wanted to determine whether the survival of neurons was compromised in this new medium when culturing cells for extended periods of time. Both human and rat primary neurons cultured in BPI for 21 days exhibited healthy cell bodies and the expected neurite networks and showed no reduction in synaptic marker Synapsin1 **(Figure 7A)**. Importantly, the total number of live human or rat neurons in culture after 21 days in BPI was no different than in standard BrainPhys medium (**Figure 7B**).

**Figure 7.**
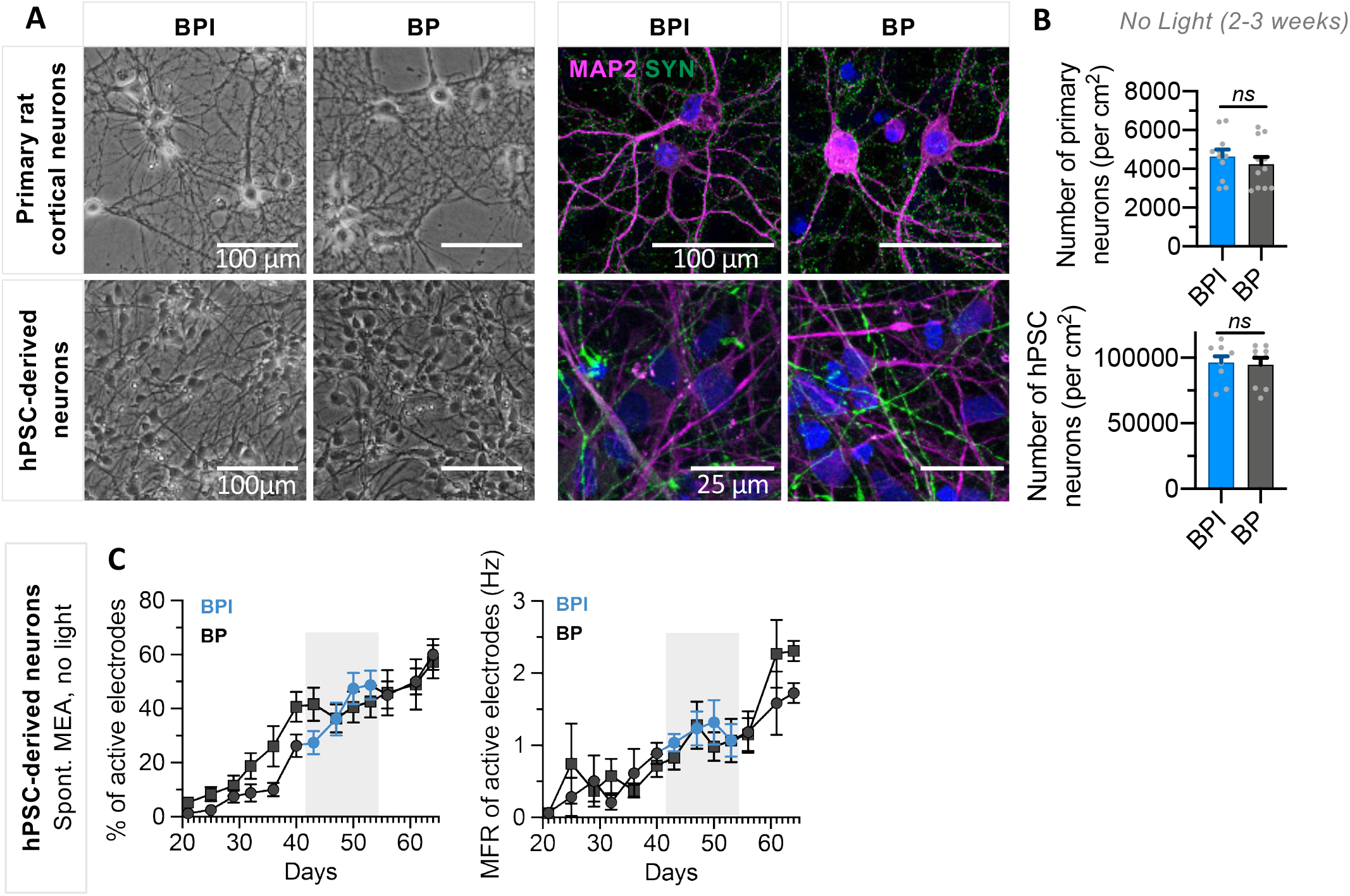
BrainPhys Imaging can support medium-term viability and function *in vitro*. **A,** shows representative images at 10x magnification showing 14 days-in-vitro (DIV) rat cortical primary neurons and hPSC-derived neurons matured in BP and BPI exhibit healthy neuron morphology. Representative immunocytochemistry images showing 21 DIV primary rat cortical neurons and 14 DIV hPSC-derived neurons characterized by MAP2 expression (magenta) matured in BP or BPI also display an appropriate expression of synaptic marker Synapsin1 (green). **B,** quantification of the number of neurons per cm^2^ shows that BPI supports equivalent neuronal survival relative to BP for 21 DIV rat cortical primary neurons (n=11) and 14-21 DIV hPSC-derived neurons (n=9). Values represent mean ± SEM. Significance determined using two-tailed non-parametric unpaired (Mann Whitney) tests. **C,** MEA recordings from hPSC-derived neurons cultured a 48-well MEA plate in BP for 9 weeks (6 wells) or switched to BPI from BP during weeks 6-8 (5 wells) represent the spontaneous mean firing rates and percentage of active electrodes (>0.017 Hz) for each data set. The percentage of active electrodes and mean firing rates were both maintained during the period in BPI. Values are shown as mean ± SEM.

To determine if human neurons remain functional for extended periods in BPI, iPSC-derived neurons were tested on an MEA. After maturation for 6 weeks in standard BrainPhys medium, half the MEA plate was switched to BPI for a period of two weeks (**Figure 7C**). Spontaneous mean firing rates and the proportion of active electrodes were similar between the wells kept in standard BrainPhys and those switched to BPI. Switching the cells back into the original BrainPhys after that period also did not affect the activity (**Figure 7C)**. Altogether, our data show that neuronal cultures are viable and functional when replacing standard BrainPhys for BPI with appropriate supplements for an extended period (2-3 weeks). For longer time periods, it is possible to alternate the feed with standard BrainPhys to replenish the light-sensitive nutrients reduced in BPI.

## Discussion

BrainPhys Imaging basal medium was designed to enhance imaging capacity and support neuronal function *in vitro*. This new medium proved advantageous for light-driven analysis of cultures of reprogrammed human brain cells and primary neurons. The most commonly used fluorophores in cell biology have a spectrum of excitation and emission from ultraviolet (~300 nm) to infrared. From ultraviolet to infrared, our data show that BrainPhys Imaging optimizes the absorbance, autofluorescence, signal-to-background ratio, and phototoxicity, and outperforms other tissue culture media. Our data also shows that BrainPhys Imaging supports physiological action potentials, synaptic activity, calcium spikes, and optogenetics function while supporting the long-term viability required for neuronal cultures.

### Cell types and neuronal models *in vitro* compatible with BrainPhys Imaging

We have shown that BPI can be used with multiple neuronal cell types *in vitro*, including human and rat, primary or reprogrammed, cortical or midbrain, 2D or 3D. Although we could not test all the possible experimental designs, we predict that most brain cell models *in vitro* or *ex-vivo* should benefit from BPI in light-driven experiments. No experiment that we performed indicated that BPI might be limited to a particular neuronal cell type. Any cell type supported by standard BrainPhys will benefit from switching temporarily to BPI during light-driven experiments. Patient-derived iPSC are increasingly popular for pre-clinical models of neurology ^28–39^. BrainPhys medium and the new imaging formula presented here were specifically designed with the intent to improve such human neuronal models.

### Live imaging technologies compatible with BrainPhys Imaging

We do not foresee any particular limitation in the type of live imaging experiment benefiting from BPI compared to the other neuronal media tested *in vitro*. The improvement in fluorescent signals and minimal photocytotoxicity in BPI can help most light-driven neuronal experiments, in particular, the ones with low signals and requiring higher light stimulation power ^19^. In addition, the optimal support of electrophysiological activity is necessary for light-driven technologies used to monitor neuronal function (e.g. voltage-sensors, calcium-sensors, synaptic imaging, pH-sensors, mitochondria motility) ^3, 5–9, 11^. The discovery of personalized treatment with patient-derived neurons will benefit from drug screening *in vitro* ^40,41^. Most drug screens use high-throughput microscopy to assess cell survival, which can be confounded by phototoxic media. Our data suggest that BPI will be the medium of choice for live-cell high-content imaging experiments in neurological and psychiatric research.

### Photo-modulation of neuronal function compatible with BrainPhys Imaging

In the last decade, optogenetics has become a major research tool to interrogate the function of nerve cells in health and disease ^3, 5, 8, 42, 43^. The integration of optogenetics with neural stem cells has enabled selective excitation and/or inhibition of specific cell populations expressing optogenes with exceptional precision, allowing researchers to investigate neural network function from innovative perspectives ^44–48^. As BPI reduces photo-toxicity and optimizes electrical and synaptic activity, such conditions certainly improve optogenetics ability to assess neural networks. We have demonstrated that BPI is highly compatible with modulating the activity of human neurons *in vitro* with light excitation of ultrafast channel rhodopsin (ChETA). Similarly, BPI may also become advantageous for other optical experiments such as the photo-release of caged compounds ^14, 15^, or optical activation of intracellular antibodies^16^.

### Long-term cultures in BrainPhys Imaging

Standard BP with appropriate supplements and tissue culture techniques is capable of sustaining neuronal maturation for over six months *in vitro* ^1^. In BPI, several components were reduced, removed or replaced from the standard BP to improve imaging capacity. We have shown that despite these modifications, BPI is capable of supporting cell viability and function (human neurons or rat primary neurons) for at least several weeks to similar levels observed in standard BrainPhys. However, BPI was designed to complement standard BrainPhys rather than replace it completely. It is hard to predict the viability effect of a medium on every cell type used in neuroscience research. It is possible that some cell types might be more sensitive than others. In some cases, it may be best to culture the cells in standard BrainPhys for the majority of the time and only switch to BPI when required for imaging assays. For longer-term imaging experiments of particularly sensitive cells, it is also possible to replenish the cells with standard BP every week or two. Despite this note of caution, based on our experiments, we are confident that using BPI for optical assays (short-term and long-term) will be better than using standard neuronal media. It is also important to note that like any other basal media, long-term viability requires the addition of appropriate supplements. In this study, we only tested the supplements generally recommended with standard BrainPhys (SM1 with or without Retinoic acid, N2-A, BDNF, GDNF, Ascorbic Acid, laminin, cAMP, IGF-I), but it may be possible to obtain similar results with different sets of supplements if they are not phototoxic.

### Advantages of BrainPhys Imaging compared to other basal media available for long-term neuronal culture and live-cell imaging

To surpass reported light-induced damage on cells cultured in Neurobasal or DMEM, two photostable basal media were developed: BrightCell™ NEUMO and FluoroBrite™ DMEM ^26^. Our analysis confirmed some improvements in the autofluorescence of these media compared to standard media (Neurobasal, DMEM/F12 or standard BP). However, both media still exhibited mild autofluorescence and increased absorbance in shorter wavelength light, whereas the autofluorescence of BPI was significantly reduced throughout a wider light spectrum (300-800nm). We do not know the cause of the residual fluorescence at shorter wavelengths in the other tissue culture media specialized for imaging, but it is likely that not all photo-active components were adjusted appropriately in these formulations. The optical superiority of BPI compared to other imaging media may be particularly prominent for experiments requiring peak excitation between 300 and 400 nm, such as blue fluorophores (e.g. DAPI, Hoechst, eBFP, Brilliant Ultraviolet, DyLight 405), calcium sensors (e.g. Fura-2), or other cellular sensors (e.g. Lysotracker Blue/MeOH). Compared to standard media such as BrainPhys, we observed significant increases in signal-to-background ratios in BPI with blue and green fluorophores. The prominence of the improvements may vary depending on the quality of the fluorophore and imaging system (e.g. lens magnification, numerical aperture values). No major differences were found in imaging red fluorophores in BPI and BP, which is likely attributed to the low absorbance and autofluorescence for long wavelengths (**Figure 1B, C**). NEUMO was specifically designed for the culture of neuronal cells and to replace the Neurobasal medium during live-cell imaging ^26^. However, when tested on human neurons *in vitro*, it failed to support spontaneous calcium spikes or optogenetic-induced action potentials. In general, the electrophysiological activity in NEUMO was substantially reduced compared to side-by-side recordings in ACSF or BrainPhys media. This result was consistent with multiple electrophysiological techniques including patch-clamping, calcium imaging and MEA on human PSC-derived neurons. It also aligns with previous data obtained comparing the electrophysiological activity of human PSC neurons in standard BrainPhys and Neurobasal ^1^. FluoroBrite DMEM is another photostable medium developed to replace DMEM in experiments requiring light stimulation. Like standard DMEM, FluoroBrite was not designed for neuronal cultures. Live calcium imaging data suggest that FluoroBrite is suboptimal for neuronal culture function. However, it is possible that it is better suited for other cell types. The suboptimal calcium spikes observed in NEUMO and FluoroBrite media may be related to their formulations driving higher calcium store release and consequently suppressing neuronal activity by activating calcium-gated potassium currents and increasing action potential thresholds^49^. In a blind comparative analysis, our data show that BPI surpasses alternative photostable basal media when used on functional human PSC-derived neuronal culture.

### Concluding remarks

Optically driven technology is known to induce phototoxic by-products in culture, leading to cellular death and metabolic alterations after light exposure. Furthermore, currently available imaging media, which were designed to limit phototoxicity, impair electrophysiological activity and fail to support fundamental neurophysiological properties. The design of BrainPhys Imaging is timely with the rapid expansion of functional imaging and human neuronal models *in vitro*. BrainPhys Imaging, which is a reformulation of the standard BrainPhys basal medium, is found to improve optical properties throughout the entire light spectrum required for fluorophores and sensors used in cellular biology. The optically optimized properties of BrainPhys Imaging will enhance the quality of results for a plethora of light-driven experiments *in vitro*. Such technical advancement provides a new useful tool for preclinical models *in vitro* and may, in turn, facilitate translation in neurology and psychiatry.

### Online Methods

All catalog numbers for reagents, antibodies, biological samples, and additional resources can be found in the *supplementary key resource table* with corresponding source details.

### Human Pluripotent Stem Cell (hPSC)-derived Neuron Culture

Human neuron cultures were derived from pluripotent stem cells in two independent laboratories using slightly different protocols.

***Method #1 used in Bardy Lab (Figures 2, 3, 4, 5, 6, S2, S3, S6-11)***: WA09 (H9) ES cell colonies (WiCell, Wisconsin, U.S.A.) were maintained in mTeSR™1 (STEMCELL Technologies) as per manufacturer’s instructions on cell culture ware coated with hESC-qualified Matrigel (Corning). The cells were split by Dispase treatment (1 U/ml; STEMCELL Technologies) and mechanical scraping every 3-5 days onto fresh Matrigel. Embryoid body-based neural induction was performed based on a previous protocol ^1, 50^ with modifications. Briefly, hESCs were cultured in an ultra-low attachment plate in a neural induction medium (NIM) consisting of DMEM/F12 with 15 mM HEPES, 1x SM1, 1x N2-A, 10 μM SB431542 (STEMCELL Technologies) and either 500 ng/ml Noggin (PeproTech) or 100 nM LDN193189 (STEMCELL Technologies). Media was exchanged every other day for 7 days to allow embryoid bodies (EBs) formation. EBs were transferred to a tissue culture dish coated with 10 μg/ml poly-L-ornithine (Sigma) and 5 μg/ml laminin (Thermo Fisher Scientific) in NIM supplemented with 1 μg/ml laminin. Media was exchanged every other day for 7 days until neural rosettes were clearly visible. Neural rosettes were manually selected under phase contrast (EVOS XL Core Imaging System, Thermo Fisher Scientific) and transferred onto fresh Matrigel-coated plates in neural progenitor medium (NPM). ***Midbrain neural progenitor differentiation***: Midbrain neural progenitor medium (mNPM) was composed of DMEM/F12+GlutaMAX (Thermo Fisher Scientific) supplemented with 1x SM1, 1x N2-A, 200 ng/ml Sonic Hedgehog (PeproTech), 100 ng/ml FGF8b (PeproTech), 200 nM ascorbic acid (Sigma) and 1 μg/ml laminin. ***Cortical neural progenitor differentiation***: Cortical neural progenitor medium (cNPM) consisted of DMEM/F12+GlutaMAX plus 1x SM1, 1x N2-A, 10 ng/ml FGF2 (STEMCELL Technologies), 200 nM ascorbic acid and 1 μg/mL laminin. Neuronal progenitor cells (NPCs) were maintained at high density, fed every other day with fresh NPM and split about once a week at 1:3-1:4 ratio onto Matrigel-coated plates using Accutase (STEMCELL Technologies). For neuronal maturation, NPCs were dissociated and seeded at a density of 8 × 10^4^ to 2 × 10^5^ cells/cm^2^ (cell line dependent) onto tissue culture plates coated with 10 μg/mL poly-L-ornithine and 5 μg/mL laminin in neural maturation medium (NMM). ***Midbrain neuronal maturation medium*** (“mNMM”) was composed of BrainPhys™ supplemented with 1x SM1, 1x N2-A, 20 ng/ml BDNF (STEMCELL Technologies), 20 ng/ml GDNF (STEMCELL Technologies), 0.5 mM dibutyryl cyclic-AMP (Sigma), 200 nM ascorbic acid and 1 μg/ml laminin. ***Cortical neuronal maturation medium*** (“cNMM”) was composed of BrainPhys™ supplemented with 1x SM1 without vitamin A, 1x N2-A, 20 ng/ml BDNF, 20 ng/ml GDNF, 10 ng/ml IGF-I (STEMCELL Technologies), 0.5 mM dibutyryl cyclic-AMP, 200 nM ascorbic acid and 1 μg/ml laminin. Half media changes with NMM were performed every 2-3 days. To improve neuronal culture quality and viability and achieve a more even distribution of cells for final neuronal maturation, post-mitotic neuronal cultures 14 days post-maturation were routinely sorted (BD FACSMelody cell sorter, BD Biosciences) and live (DAPI negative) cells were replated at a density of 1.6 to 2.1 × 10^5^ cells/cm^2^ (cell line dependent) on poly-L-ornithine- and laminin-coated glass coverslips or tissue culture-treated plates in NMM containing half concentration of growth factors.

***Method #2 used in SCT Lab (Figure 7A, B)***: Frozen neural progenitor cells (NPCs) derived from embryonic (ES) or induced pluripotent stem (iPS) cells (previously characterized in ^51^) were differentiated into neuronal precursors with the STEMdiff™ Neuron Differentiation Kit (STEMCELL Technologies). Neuronal precursor cells were then frozen with Cryostor CS10 (STEMCELL Technologies) in liquid nitrogen storage and thawed later for experiments. Post-thaw, cells were plated in 500 μL per well of STEMdiff™ Neuron Differentiation medium at 26,000 to 130,000 cells per cm^2^ (cell line dependent) in a 24-well plate. On day 1 post-thaw, 500 μL of BrainPhys™ or BrainPhys™ Imaging supplemented with 1x NeuroCult™ SM1 (STEMCELL Technologies), 1x N2-A (STEMCELL Technologies), 20 ng/ml BDNF (STEMCELL Technologies), 20 ng/ml GDNF (STEMCELL Technologies), 200 nM ascorbic acid (STEMCELL Technologies), 1 μg/ml laminin (Sigma) and 0.5 mM dibutyryl cyclic-AMP (STEMCELL Technologies) [complete BrainPhys™] was added per well. Half media changes were performed every 3-4 days with complete BrainPhys™ or BPI.

### Neuronal Basal Media Tested

ACSF contained (in mM) 121 NaCl, 4.2 KCl, 1.1 CaCl_2_, 1 MgSO_4_ (or 0.4 MgSO_4_ and 0.3 MgCl), 29 NaHCO_3_, 0.45 NaH_2_PO_4_-H_2_O, 0.5 Na_2_HPO_4_ and 20 glucose (all chemicals from Sigma). BP, BPI, and BP without phenol red were obtained from STEMCELL Technologies. FluoroBrite, DMEM/F12, and Neurobasal were obtained from Thermo Fisher Scientific, NEUMO was obtained from Merck. For long-term maturation of human neuronal culture, all basal media were supplemented with: ***Human midbrain neurons supplements:*** 1x N2-Supplement A, 1x NeuroCult™ SM1, 200 nM ascorbic acid, 20 ng/mL or 10 ng/mL BDNF, 20 ng/mL or 10 ng/mL GDNF (STEMCELL Technologies), 1 μg/mL laminin (Thermo Fisher Scientific), and 0.5 mM or 0.25 mM dibutyryl cyclic-AMP (Sigma). ***Human cortical neurons supplements:*** 1x N2-Supplement A, 1x NeuroCult™ SM1 without vitamin A, 200 nM ascorbic acid, 20 ng/mL or 10 ng/mL BDNF, 20 ng/mL or 10 ng/mL GDNF, 10 ng/mL or 5 ng/mL IGF-I (STEMCELL Technologies), 1 μg/mL laminin (Thermo Fisher Scientific), and 0.5 mM or 0.25 mM dibutyryl cyclic-AMP (Sigma). Two concentrations are listed for growth factors and dibutyryl cyclic AMP since these factors were reduced by 50% in the media after two weeks of neuronal maturation.

### Dorsal Forebrain Organoid Culture

WA01 (H1) embryonic stem cells (WiCell) were cultured in mTeSR™1 as per the manufacturer’s instructions (MA29106, STEMCELL Technologies) plated on Matrigel™ (Corning). A stable GFP overexpressing cell line was generated by transfecting cells with a plasmid encoding GFP expression driven by the chicken beta-actin promoter ^52, 53^ using the Neon Transfection System (Thermo Fisher Scientific). H1 cells were dissociated to single cells and 1.2×10^6^ cells were resuspended in 120 μL of R-buffer containing 2μg of pCAG:GFP:IRES:Puro. Cells were then electroporated using the manufacturer’s instructions with the following settings: 1,200 volts, 20 ms, 1 pulse. The transfected cell suspension was then added to 4 mL of mTeSR™1 supplemented with CloneR™ (STEMCELL Technologies), mixed by pipetting, then added to tissue culture-treated plates coated with Matrigel™ at 2.5 ×10^4^ cells/cm^2^. The medium was replaced after 24 hours and fed with fresh medium daily until the cells were ~60% confluent. The cells were then fed with fresh medium containing 1μg/mL puromycin (Cayman Chemicals) daily to select for stable transfectants. Clonal cell lines were then generated by dissociating the H1-GFP cells to single cells and seeding at 10 cells/cm^2^ in mTeSR™1 supplemented with CloneR™ onto tissue culture-treated plates coated with Matrigel™. Cells were then fed 2 days post-plating with fresh medium supplemented with CloneR™, then fed daily following 4 days post-plating with mTeSR™1. Colonies were then picked manually between 7-9 days and cultured as above. Dorsal forebrain organoids were generated as previously described ^54^ with the following modifications; hPSCs were single-cell dissociated using Gentle Cell Dissociation Reagent (STEMCELL Technologies) for 7 minutes at 37⁰C. To generate ventral forebrain organoids, H1-GFP forebrain organoids were treated with IWP-2 (5μM, Selleckchem) and SAG (100nM, Selleckchem) between day 6 to 25 of the dorsal forebrain organoid protocol.

### Primary Rat Neuron Culture (Figures 2E-F, 3A-D, 3K, 7A-B, S1C-F, S4B,C, S5)

Pairs of Rat E18 cortices were obtained from BrainBits, LLC and dissociated for 10 minutes in papain (at least 20U/ml; Worthington Biochemical). A single-cell suspension was obtained and filtered through a 40μm cell strainer. The resulting primary cells were cultured in NeuroCult™ Neuronal Plating medium (STEMCELL Technologies) or Neurobasal™ medium (for Neurobasal and NEUMO cultures) with 1x SM1, 0.5mM L-glutamine and 25μM L-glutamic acid on culture-ware pre-coated with 10μg/mL poly-D-lysine (Sigma). Cells were plated at 30,000 cells/cm^2^ in a 24-well plate. 5 days post-plating, half media changes were performed every 3-4 days with either BrainPhys™ (STEMCELL Technologies), Neurobasal™, BrightCell™ NEUMO (Merck) BrainPhys™ Without Phenol Red (STEMCELL Technologies), or BrainPhys™ Imaging (STEMCELL Technologies) supplemented with 1x SM1. Neurobasal™ and BrightCell™ NEUMO media were also supplemented with 0.5mM L-glutamine.

### Osmolality Measurements

To assess media osmolality, 20μL of test media or supernatant collected from maturing human iPSC-derived cortical neurons before feeding was pipetted into a disposable tube (Advanced Instruments). A Fiske Micro-Osmometer (model 210) was used and calibrated with 2000, 850, 290 and 50 mOsmol/kg standards. Before recordings, the calibration was referenced against 290 mOsmol/kg standards, and between recordings, the micro-osmometer probe was cleaned with a probe cleaner (Advanced Instruments). Three human cerebrospinal fluid (hCSF) samples were pooled from multiple subjects (1 to 4).

### Absorption Spectra of Culture Media

Absorption spectra were measured using a CARY 7000 spectrophotometer (Agilent Technologies, Santa, CA, USA) with an integrating sphere attachment and a cuvette center mount holder designed to measure the absorbance of liquid samples ^55^. A 1 cm pathlength cuvette with all four clear sides was used with 3 ml of sample for each measurement and spectra were measured from 200-800 nm in 5 nm increments with a spectral bandwidth of 2 nm. Acquired spectra were baseline corrected and blank subtracted to obtain corrected absorption spectra.

### Autofluorescence Measurements of Culture Media

Media autofluorescence was tested in two independent laboratories using slightly different protocols and apparatus.

***Method #1 (figure 1C):*** Volumes (10 mL) of control and test media were aliquoted into a blacked-out 15 mL tube (Corning). A custom bifurcated fiber probe (200 μm 0.22 NA) was dipped into the media. 375nm, 405nm, 488nm, 532nm excitation wavelengths (20 mW) were sequentially fired from a Toptica iCHROME MLE (Farmington, NY, USA). Emission spectra (400 nm – 700nm) from each excitation wavelength were captured using a Horiba MicroHR (Kyoto, Kyoto, Japan) using a 500 ms integration time. The following Semrock (Rochester, NY, USA) Razoredge long-pass filters were used for 375nm & 405nm, 488nm, 532nm generated spectra respectively: 405nm, 488nm, 532nm. ***Method #2 (figure 1D-F):*** Equal volumes (300 μL) of control and test media were pipetted into a CellCarrier-96 black plate with an optically clear bottom (Perkin Elmer). A total of eight replicate wells per medium were tested with a BMG Labtech FLUOstar Omega using an excitation/emission of 355/460 (blue), 485/520 (green), 544/590 (red) and 584/620 nm (far-red). FLUOstar uses a high energy xenon flash lamp light source with a 100 Hz flash rate and 167 ms exposure time. The energy per flash emitted is 67 mJ and subjects each well to 1mW of light power. Emission wavelengths were captured using a BMG Labtech photomultiplier tube detector (PMT) at a gain of 1000. Blue, green, red and far-red filters utilized the respective bandwidths for their excitation/emission wavelengths: 20/20, 12/25, 20/20 and 10/20 nm. Acquired autofluorescence data was blank subtracted. For normalization, the mean fluorescence intensity in PBS was subtracted from the other media.

### Lentiviral Vectors

Lentiviral vectors were produced in Lenti-X™ 293T cells (Takara Bio) cultured in DMEM, high glucose, no glutamine (Thermo Fisher Scientific) supplemented with 10% fetal bovine serum (Thermo Fisher Scientific), 4 mM GlutaMAX supplement (Thermo Fisher Scientific) and 1 mM sodium pyruvate (Sigma) on cell culture-ware coated with 0.002% poly-L-lysine (Sigma). Lenti-X™ 293T cells were transfected with 12.2 μg lentiviral transfer plasmid (LV Syn-hChR2-T159C-E123T-EYFP-WPRE [hSyn:ChETA-YFP; ^12^], or LV pCSC-Synapsin(0.5kb)-MCS-EGFP, a derivative of pCSC-SP-PW-EGFP) and packaging plasmids (8.1 μg pMDL/RRE, 3.1 μg pRSV/REV and 4.1 μg pCMV-VSVg) using a polyethylenimine (PEI, 25 kDa, linear; Polysciences) transfection method (4:1 ratio of PEI to DNA). The culture medium was exchanged six hours after transfection. The supernatant containing lentiviral particles was collected ~66 hours after transfection, filtered through a sterile 0.45-μm SFCA membrane filter (Corning) and ultracentrifuged at 25,000 rpm (Beckman) at 4°C for 2 hours. Virus pellets were resuspended in Hank’s Balanced Salt Solution (HBSS; Thermo Fisher Scientific) and virus titers were determined using the Lenti-X™ qRT-PCR Titration Kit (Clontech) according to the manufacturer’s instructions. Neuron cultures were transduced with titer-matched lentiviral vectors (2.67×10^3^ viral RNA copies/cell) for 48 hours prior to wash. Lentiviral transfections in neuronal cultures were performed at least one week prior to experimentations. If expression levels were low, neurons were left in culture for an extra 3-5 days before further experimentations commenced.

### Immunofluorescent Staining Protocol

Cultures were fixed in 4% paraformaldehyde (Alfa Aesar) at 22°C for 30 minutes. Cells were then permeabilized with 0.1% Tween-20 (Sigma) in PBS (PTW) for 15 minutes. Cells were incubated overnight at 4°C with anti-β-III-tubulin antibody (Biolegend; 1:1000), anti-MAP2 antibody (Abcam; 1:5000) and/or anti-Synapsin I antibody (Merck AB1543; 1:2000) diluted in PBS containing 10% normal donkey serum (Merck). The next day cells were washed three times with PTW and incubated with anti-mouse Alexa Fluor®488 (Jackson Immunoresearch; 0.625μg/mL), anti-rabbit Alexa Fluor®488 (Jackson Immunoresearch; 0.625μg/mL), anti-chicken Alexa Fluor®488 (Jackson Immunoresearch; 0.625μg/mL), anti-chicken Alexa Fluor®647 (Jackson Immunoresearch; 0.625μg/mL) and/or anti-mouse DyLight®594 (Thermo Fisher Scientific; 1 μg/mL) diluted in PBS containing 2% normal donkey serum overnight at 4°C. Cells were then washed three times with PTW to remove residual antibodies. To detect the cell bodies, Hoechst 33342 (Sigma, 2μg/mL) was used to counterstain the cells.

### Neurofluor™ NeuO Labelling Protocol

E18 rat cortical neurons were cultured in BrainPhys™, BrainPhys™ Imaging or BrainPhys™ without phenol red for 11 days (16 days total) before labeling with NeuroFluor™ NeuO (STEMCELL Technologies). For labeling, the culture medium was aspirated and replaced with a labeling medium consisting of culture medium plus 0.25μM NeuroFluor™ NeuO. Cells were incubated in the labeling medium for 1 hour at 37⁰C. After incubation, the labeling medium was replaced with fresh medium and cells were incubated at 37⁰C for 2 hours before imaging.

### Image Acquisition

Either an SP8 confocal microscope (Leica), ImageXpress Micro 4 High Content Screening System (Molecular Devices) with an Andor SDK3 camera, CKX53 inverted microscope with an SC100 digital camera (Olympus), or a BX51 upright microscope (Olympus) with a PCO.Panda 4.2 digital camera were used for image acquisition. All images were taken using 16-bit cameras. Varying lens parameters and acquisition settings were used, as listed below.

### Imaging Fixed Primary Rat Cortical Neurons (Figure 2E, 7A,B and S1C, E, F)

Fixed primary rat cortical neurons were acquired using an SP8 confocal microscope or an ImageXpress Micro 4 High Content Screening System. In figure 2E and S1E, F, β-III-tubulin (TexasRed filter set: 562±20/624±20 nm, dichroic filter 593nm) was imaged using the ImageXpress Micro 4 High Content Screening System at 500 ms with a 20x air immersion lens (Ph1 S Plan Fluor ELWD ADM; NA 0.45). For fluorophore excitation, the in-built Lumencor Sola Light (SE 5-LCR-QB) was set to 100 “lumencor intensity” units in MetaXpress (v6.2.2.) software. This same set-up was used in figure S1C for Hoechst (DAPI filter set: 377±25/447±30 nm, dichroic filter 409 nm) with a 204ms exposure time and figure 7B for β-III-tubulin (FITC filter set: 475±17/536±20 nm, dichroic filter 506 nm) and Hoechst (DAPI filter set) with 100ms and 204ms exposure times respectively. For figure 7A and B, Synapsin I, MAP2 and Hoescht were imaged using the SP8 confocal microscope with a 63x oil objective (HC PL APO CS2; NA 1.4) and the following settings: 405nm, 488nm and 638nm lasers with 26.7%, 18.3% and 1% respective power intensities. The following detector settings for Synapsin I, MAP2 and Hoescht images were respectively selected for: 410nm − 483nm (HyD) at gain 100, 493nm - 643nm (PMT) at gain 768.8 and 643nm - 781nm (HyD) at 40.3, Pinhole 1AU, zoom 1x, and 400Hz scan speed.

### Imaging Fixed hPSC-derived Neurons (Figure 7A, B)

Images of fixed hPSC-derived neurons were acquired using a Leica SP8 confocal microscope or ImageXpress Micro 4 High Content Screening System. MAP2 (Cy5 filter set: 628±20/692±20, dichroic filter: 660 nm) and Hoechst (DAPI filter set) from figure 7B were both imaged using the ImageXpress Micro 4 High Content Screening System with a respective 50 ms and 204ms exposure time using a 20x air immersion lens (Ph1 S Plan Fluor ELWD ADM; NA 0.45). For all image acquisition using the ImageXpress, the in-built Lumencor Sola Light (SE 5-LCR-QB) was set to 100 “lumencor intensity” units in MetaXpress (v6.2.2.) software. In figure 7A, Synapsin I, MAP2 and Hoechst were imaged using a Leica SP8 confocal microscope with a 63x oil objective (HC PL APO CS2; NA 1.4) and the following settings: 405nm, 488nm and 638nm lasers at 1%, detectors 410nm - 483nm (HyD) at gain 142, 493nm - 643nm (PMT) at gain 768.8 and 643nm - 781nm (HyD) at 32.5, Pinhole 1AU, Zoom 1x, Scan speed 400 Hz.

### Imaging Live Primary Rat Cortical Neurons (Figure 2E, 3A-C, S1D, F, S4B, S5, 7A)

Live rat cortical neurons labeled with Neurofluor™ NeuO (FITC filter set: 475±17/536±20 nm, dichroic filter 506 nm) and Hoechst (DAPI filter set: 377±25/447±30 nm, dichroic filter 409 nm) were imaged using an ImageXpress Micro 4 High Content Screening System set at 25ms and 204ms exposure times respectively with a 20x air immersion lens (Ph1 S Plan Fluor ELWD ADM; NA 0.45). NeuroFluor™ NeuO has an excitation/emission spectrum of 470/555nm. The in-built Lumencor Sola Light (SE 5-LCR-QB) was set to 100 “lumencor intensity” units in MetaXpress (v6.2.2.) software for fluorophore excitation. Phase-contrast images (Figures 3A-C, S4B, S5, 7A) of live rat cortical neurons were taken with an Olympus CKX53 microscope with an SC100 digital camera and a 10x dry objective (CACHN10XIPC; NA 0.25).

### Imaging Live Fused Dorsal and Ventral Forebrain Organoids (Figure 2D)

Fused dorsal and ventral forebrain organoids were imaged using a Leica SP8 confocal microscope with 10x dry objective (HC PL APO CS2, NA 0.4) and the following settings: 488nm laser at 5% power intensity, detector 493nm - 547nm (HyD) at gain 176.8, Pinhole 1AU, Zoom 1x, Scan speed 600 Hz.

### Imaging Live hPSC-derived Neurons (Figures 2A-C, S2A-B, S3, 7A)

Fluorescent images of live hPSC-derived neurons transfected with green-fluorescent-protein (GFP) lentivector were imaged at 200ms exposure time using a PCO.Panda 4.2 (16-bit) digital camera and Olympus BX51 upright microscope with 40x water immersion lens (LUMPLFLN40XW, 0.8NA, Semiapochromat). A cool-LED pE300 illumination unit was set to blue (460 nm) at 1% power (0.1 mW) and light passed through a single band FITC/Cy2 filter (Chroma, 49002) with a 470 nm excitation (±40 nm) and 525 nm emission (±50 nm) wavelength. Single neurons were perfused with BrainPhys™ basal media bubbled with a mixture of CO_2_ (5%) and O_2_ (95%) and maintained at room temperature. Perfusates were then switched to BrainPhys™ Imaging and then ACSF. Imaging parameters remained constant across all live neuron images in each perfusate. Phase-contrast images (Figure 7A) of live hPSC-derived neurons were taken with an Olympus CKX53 microscope with an SC100 digital camera and a 10x dry objective (CACHN10XIPC, NA 0.25).

### Quantification of Cell Viability with Imaging for Phototoxicity Experiments

***Live Primary Rat Cortical Neurons (Figure 3D, 7B):*** Neuron cultures for quantification were set up in triplicate wells and fixed for immunostaining with β-III-tubulin. Twenty-five images per well were taken using an ImageXpress Micro 4 High Content Screening System (Molecular Devices) for a total of 75 images per condition. Using a custom module designed for neuron quantification in MetaXpress 6.2.2 software (Molecular Devices), neurons in each image were counted based on the following parameters. Nuclei are masked if between 10.5 and 35 μm in width and 90 and 350 μm2 in area with an intensity above the local background of 500. Objects with a max intensity standard deviation above 2000 were excluded (excludes apoptotic nuclei). Masked area is then queried for presence of β-III-tubulin positive staining with an average intensity above 400. Images, where the neuron count was above 55, were excluded due to increased error in the presence of high nuclei counts. The average number of neurons per cm^2^ was calculated based on the image size (0.00508cm^2^) produced from the 20x objective. **Fixed hPSC-derived Neurons (Figure *7B*)**: Neurons for quantification were set up in duplicate wells and fixed for immunostaining on days 14-21. Sixteen images were taken per well and quantified as described above based on the presence of MAP2 positive staining. The average number of neurons per cm^2^ was calculated as described above.

### Fluorescent Signal-to-Background Ratios

***iPSC-derived Human Neurons (Figures 2B, C)***: The mean ‘signal’ intensity of human neurons expressing synapsin:GFP were analyzed per field-of-view (FOV) by computing the mean gray values of regions of interest (ROI) manually selected around soma and/or neurites with ImageJ’s ‘multi-measure’ function ^56^. Whether the signal intensity at soma and neurites were analyzed together or individually is detailed in figure legends. To analyze the mean ‘background’ intensity in each test media, ROIs were traced around regions without neurons (i.e. cellular bodies or neurite projections) and the mean gray values generated. Signal-to-background ratios were calculated using the following formula: ((signal intensity – background intensity)/background intensity). Per FOV, ‘signal’ and ‘background’ analyses were repeated using the same ROIs across test conditions. Imaging parameters of images quantified are included under *‘Imaging Live hPSC-derived Neurons’* (GFP). ***Primary Rat Cortical Neurons (Figures 2F, S1C-E):*** <u>Mean ‘signal’ intensity</u> analysis on the fluorescence staining of fixed and live primary rat cortical neurons was conducted per FOV using ImageJ’s ^56^ ‘3D object counter’ (Figure 2F) or ‘Triangle’ auto-threshold (Figure S1C-E) in mean gray value units. In figure 2F, the fluorescence ‘signal’ from Hoechst, NeuO and β-III-tubulin above 3600, 600 and 7000 (mean gray value units) were masked, respectively. Mean ‘signal’ intensity values of masked regions greater than 30 pixels in size (pixels^2^) were computed with ‘3D object counter’. In figures S1C-E, ROIs of Hoechst, NeuO and β-III-tubulin fluorescence stains were generated from binary masks obtained from auto-selection/thresholding. ROI outlines were overlapped onto the corresponding original image(s). ImageJ’s ‘multi-measure’ function was used to calculate the mean ‘signal’ intensity of masked regions greater than 30 pixels in size (pixels^2^). <u>*For mean ‘background’ intensity*</u> analysis, ‘background’ ROIs (i.e. with no cellular bodies or neurite projections) were generated per FOV from binary masks obtained using ImageJ’s ‘Huang dark’ threshold (Figure 2F) or ‘Triangle dark’ auto-threshold (figure S1C-E). ‘Background’ ROIs were overlapped onto the corresponding original image(s) and the mean gray value units computed per FOV using ImageJ’s ‘multi-measure’ function ^56^. Threshold settings remained constant for each fluorescent stain to maintain consistency between replicates. Objects on edges were excluded from all mean intensity results. Signal-to-background ratios were calculated using the following formula: ((signal intensity – background intensity)/background intensity). Imaging parameters of quantified images are included under *‘Imaging Live Primary Rat Cortical Neurons’ and ‘Imaging Fixed Primary Rat Cortical Neurons’*.

### Image Analysis Per Pixel (Figure 2B, S2B)

GFP intensity values were computed per pixel from ROIs selected at ‘background’, 'neurites' and 'soma' regions, as described above (‘Fluorescent Signal-to-Background Ratios’ for *iPSC-derived neurons*). Using Image J’s ‘Histogram’ command ^56^, ROIs were analyzed using 1240 bins in the intensity range from 60 to 1300. Mean GFP intensity results across all FOVs were graphed either against the cumulative percentage of pixels (Figure 2B) or the number of pixels (figure S2B). All data were paired across test conditions. Imaging parameters of GFP images quantified are included under *‘Imaging Live hPSC-derived Neurons’.* The number of photo-counts of the camera sensor without cells and media were included and labeled as ‘dark’ (Figure 2B and S2B).

### LED Exposure for Phototoxicity Experiments (Figure 3A-D)

A string of battery-operated blue (ER CHEN), red (ER CHEN) and violet (AMARS) LED lights were wrapped around a tissue culture plate. The plate containing the lights was then placed above the test culture plate (containing cultures of primary E18 rat cortical neurons) in a temperature-controlled incubator. The test plate was exposed to light for 12 hours total (two bouts of 6 hours, 24 hours apart) for blue LEDs (450-475 nm; 14 Lux ± 1.41 S.E.M), 18 hours for red LEDs (620-740 nm; 55 Lux ± 4.99 S.E.M) and 6 hours for violet LEDs (395-405 nm; 430 Lux ± 3.70 S.E.M). Light meter readings were taken of the plate containing the LEDs at 8 points over the plate area with a Sper Scientific™ model 840020 light meter.

### Mitochondrial Membrane Potential and Cell Viability Assessment (Figures 3K, S4C)

Primary rat cortical neurons were cultured as described in section “Primary Rat Neuron Culture” in 96 well plates pre-coated with 30μg/mL poly-D-lysine (Sigma) at 23,400 cells/cm^2^. On day 13 or 14, cultures were exposed to violet LED light as in ‘*LED Exposure for Phototoxicity Experiments’* for 1 hour in a humidified incubator at 37 °C with 5% CO_2_ and 21% O_2_ with a rest period of 2-3 hours post-exposure before fluorescent assays. For H_2_O_2_ exposure, 3% (0.88 M) hydrogen peroxide solution (Sigma) was diluted to 100μM final concentration in culture wells and cells were incubated for 4 hours in a humidified incubator at 37 °C with 5% CO_2_ and 21% O_2_. For fluorescent assays, the culture medium was removed from all wells and replaced with 200μL BPI. 20μL of CellTiter-Blue® Reagent (Promega) was added per well in triplicate wells for each medium condition (BP or BPI). After incubation in 37 °C with 5% CO_2_ and 21% O_2_ for 30 minutes, JC-1 probe (Invitrogen; reconstituted in dimethyl sulfoxide) was added to the second set of triplicate wells to 2μg/mL final concentration. Plates were then incubated for another 30 minutes at 37 °C with 5% CO_2_ and 21% O_2_. Wells containing JC-1 probe were subsequently washed by performing two half-medium changes with BPI. Fluorescence was read using a SpectraMax M5 Multi-mode Microplate Reader (Molecular Devices) at wavelengths: 485nmEx/535nmEm (JC-1 monomers), 535nmEx/595nmEm (JC-1 aggregates), and 560nmEx/590nmEm (CellTiter-Blue® cell viability). Fluorescence readings were taken in triplicate for each well in a horizontal scan pattern. Blank values (BPI) were subtracted from all readings in SoftMax Pro software (version 7.1). Values were averaged across triplicate readings and wells for each condition. Ratio of JC-1 aggregates to monomers was calculated by dividing the value of 595nm emission by the value of 535nm emission.

### Media Cytotoxicity Comparisons (Figure 3E-F)

***LDH Assay for Cell Viability***: Lactate Dehydrogenase (LDH) release in human neuronal cultures treated with light-stimulated neuronal maturation media was measured to assess loss of cell viability. Media was made and stimulated with ambient light for 24 hours as described in *‘H_2_O_2_ measurements’*. Human midbrain and cortical neurons were matured and treated in 48-(Figure 3G) and 96-well (Figure 3H) culture plates, respectively. ***Treatment:*** Neuronal maturation media was replaced with 24-hour light stimulated media (Day 0). Half-Media changes were performed every 24 hours (Day 1, Day 2…) using fresh 24-hour light stimulated media. Supernatant samples were collected on Day 0 immediately *after* feeding, and immediately *before* feeding on Day 1 onwards for the LDH assay. Samples were frozen at −20°C prior to LDH measurement. ***LDH Assay:*** LDH was measured using CytoTox 96® Non-Radioactive Cytotoxicity Assay (Promega) as per the manufacturer's protocol. Formazan absorbance at 490nm was measured by GloMax microplate reader (Promega) using GloMax (v3.1) software. Blank LDH levels were subtracted from Treatment LDH values. Normalization can be found in figure legends. For Figures 3E and F, luminescence measurements were normalized first to BPI results for each respective day, then values were renormalized to day 0.

### H_2_O_2_ Measurements in Light Stimulated Media (Figure 3G, J)

**Media**: BrainPhys (BP) and BrainPhys Imaging (BPI) basal media were supplemented to make neuronal maturation media as described in ‘*Human Pluripotent Stem Cell (hPSC)-derived Neuron Culture’ (method#1 Bardy Lab*). Media was made fresh for each experiment and stimulated with blue or ambient light for 24-hours. ***Lumos optical stimulation (at 475 nm):*** 500uL of media was added to the wells of a 48-well Lumos MEA plate (Axion Biosystems). The 48-well Lumos MEA plate was maintained at 37°C, 5% CO_2_ environment within a Maestro Pro MEA system (Axion Biosystems). The media were stimulated with blue light (475 nm) for 24-hours using a Lumos optical stimulator (Axion Biosystems) adapted for 48-well MEA plates. Stimulation was repeated with the following settings: 10 × 5 ms flashes of blue LED light at 10 Hz and a 30-second inter-burst interval. Incremental blue LED power intensities were used (0, 3, 5, 12, 27, 53, 79 and 105 mW). The light intensity (mW) of blue light emitted from a Lumos optical stimulator (Axion Biosystems) was measured using a PM100D Compact Power and Energy Meter Console (Thor Labs) set to 475 nm (Figure S4A). ***Ambient light stimulation:*** 500μL of media was added to the wells of a 48-well plate (Corning CoStar). 48-well plates were placed in a Cellgard ES ClassII biological safety cabinet (Nuaire) and stimulated for 24-hours with a 58-watt NL-T8 Fluorescent Lamp Spectralux©Plus (Radium) located at the ceiling of the cabinet. The lamp is 1.5 meters in length with a 2.5 mg mercury content and medium bi-pin G13 bas, giving rise to a 5200lm luminous flux. Control plates were wrapped with alfoil to protect them from light. ***H_2_O_2_ Luminescence Measurements:*** H_2_O_2_ measurements were made using ROS-Glo™ (Promega) kits as per the manufacturer's protocol. Upon concluding light stimulation, H_2_O_2_ substrate was added to sample media to a final concentration of 25μM. Samples were incubated for 2 hours in a dark humidified incubator at 37 °C with 5% CO_2_ and 21% O_2_. Samples were transferred to an opaque black-walled 96 well plate in triplicates, combined with an equal volume of ROS-Glo™ Detection Solution, and incubated for 20 minutes before relative luminescence units (RLU) was measured on a GloMax microplate reader (Promega) using GloMax (v3.1) software.

### Light spectra recordings

***Ambient light (Figures 3H):*** To assess emission spectra from a 58-watt NL-T8 Fluorescent Lamp Spectralux©Plus (Radium) in a Cellgard ES ClassII biological safety cabinet (Nuaire), a Flame VIS-NIR spectrometer (Ocean Insight, Rochester, NY, USA) with a 10 μm slit was placed in the cabinet, facing the white-light fluorescent lamp. The emission spectra were captured using an integration time of 2ms. ***LUMOS optical stimulator (at 475nm) (Figures 3I):*** To assess emission spectra from the Lumos optical stimulator (Axion Biosystems) set to a wavelength of 475nm, a Flame VIS-NIR spectrometer (Ocean Insight, Rochester, NY, USA) installed with a 10 μm slit was placed approximately 1cm away from a single LED on the Lumos optical stimulator. The emission spectra were captured using an integration time of 1ms.

### Calcium Imaging

hPSC-derived neurons attached onto coverslips in 48-well plates were incubated with a 0.5 μL of Fluo 4-AM (1mM, Thermo Fisher Scientific) in 500 μL of neuromedium, per well (final Fluo 4-AM concentration in media: 1μM). Incubation occurred for 20 minutes in a humidified incubator at 37 °C with 5% CO_2_ and 21% O_2_. Excess dye was removed by washing three times with ACSF. The cells were then transferred into a room-temperature recording chamber continuously perfused with ACSF and bubbled with a mixture of CO_2_ (5%) and O_2_ (95%) for an additional 30 minutes at room temperature to allow de-esterification. Fluo-4 AM concentrations lower than 2μM and loading times less than 30 minutes did not induce harmful effects to the neurons ^57^. Time-lapse image sequences were acquired at 5 Hz over 4 minutes with a region of 248 × 248 pixels. Images were recorded at 200ms exposure time using a PCO.Panda 4.2 digital camera and an Olympus BX51 upright microscope with a 10x water immersion lens (UMPLFN10XW, NA0.3, Semi-Plan Apochromat) and FITC/Cy2 filter set (Chroma). A cool-LED pE300 illumination unit was set to 460 nm (1%; 0.1 mW) to excite Fluo 4-AM. Imaging parameters were kept constant when recording in different test media. To assess network activity in response to different media, the following were perfused into the bath over 6 minutes at a constant rate of 0.32 mL/min: ACSF, BPI, BP, NEUMO, FluoroBrite. At the end of the experiment before discarding the culture 1 μM Tetrodotoxin (TTX, Abcam) perfusate was added to the basal media to block voltage-gated sodium channels and action potential generation. The orders of media perfusion were alternated, apart for TTX, which was always perfused last. Images were then processed using ImageJ software. A heat-map LUT was applied to each image sequence to visualize calcium events as changes in fluorescence over baseline. A region of interest (ROI) was placed around each soma displaying a defined cellular morphology and at least one clear neuronal calcium event (dF/F >5%, fast rise, slower decay). A time series for each ROI was calculated using ImageJ ^56^ and Microsoft Excel before analysis in Clampfit (v10.7). Different types of calcium events were manually categorized as either calcium waves, spikes or a combination of calcium waves and spikes **(Figure S8A)**.

### Patch-clamping

Individual coverslips containing neurons were transferred into a recording chamber at room temperature (21 °C) continually perfused at 0.32 ml/min with either ACSF or BrainPhys™ Imaging media bubbled with a mixture of CO_2_ (5%) and O_2_ (95%). Whole patch recordings were made on neurons expressing the previously infected synapsin:GFP lentiviral vector. Targeting whole-cell recordings were achieved via a 40x water-immersion objective, and a PCO.Panda 4.2 digital camera on an Olympus BX51 microscope. A cool-LED pE300 illumination unit at 460 nm was used to visualize neurons expressing synapsin:GFP. Patch electrodes were filled with internal solutions containing 130 mM K--gluconate, 6 mM KCl, 4 mM NaCl, 10mM Na-HEPES, 0.2 mM K--EGTA, 0.3mM GTP, 2mM Mg-ATP, 0.2 mM cAMP, 10mM D--glucose, 0.15% biocytin and 0.06% rhodamine for somatic (open tip resistance 3-5 MΩ) whole-cell recordings. The pH and osmolarity of the internal solution were close to physiological conditions (pH 7.3, 290–300mOsmol). AMPA-receptor mediated events were observed exclusively in voltage-clamp at −70 mV (close to anion reversal potential). GABAa-receptor-mediated kinetics were observed exclusively in voltage-clamp at 0 mV (close to cation reversal potential). Recorded membrane potential values were adjusted for pipette offset. Electrode capacitance was compensated for in cell-attached mode. Signals were low-pass filtered (DC to 20 kHz) and acquired by PClamp software. Whole-cell recordings were amplified via a Digidata 1440A/Multiclamp series 700B. On average the neurons recorded had an access resistance of ~22+7 MΩ (Mean+Stdev). The average patch-pipette resistance used was 3.8+0.5 (Mean+Stdev). Final data were analyzed using a combination of Clampfit (v10.7) and GraphPad Prism 8 and were corrected for liquid junction potentials (10 mV).

### Action Potential (AP) Type Classification

The classification was based on previous work (Bardy et al 2016). “Type 0 cells” are most likely non-neuronal and do not express voltage-dependent sodium currents. “Type 1 neurons” express small Nav currents but are not able to fire APs above −10 mV. The limit of −10 mV was chosen as it is close to the reversal potential of cations (0 mV), and a sign of healthy mature APs. “Type 2 neurons” fire only one AP above −10 mV, which is typically followed by a plateau. “Type 3 neurons” also fire an AP above −10 mV and one or a few aborted spikes below −10 mV. “Type 4 neurons” fire more than one AP above −10 mV but at a frequency below 10 Hz. “Type 5 neurons” fire APs above −10 mV at 10 Hz or more.

### Optogenetics During Patch-clamping

Blue light stimuli (475 nm) were controlled by a computer-controlled trigger signal to the LED illuminator (CoolLED, pE300) synchronized with the pClamp protocols through a Master 9 programmable pulse stimulator (A.M.P.I). The patched neurons were stimulated with a burst consisting of 10 flashes of blue (475 nm) light of a duration of 5 ms at 10Hz. Each burst stimulus was repeated at least ten times every 10s for each cell. The intensity of the blue light at 475 nm tested here ranged between 0.1, 0.2, and 0.4 mW. Illumination was achieved via an Olympus BX51 upright microscope, through a 40x water immersion lens (LUMPLFLN40XW, 0.8NA, Semiapochromat) focused on the soma and surrounding of the patch-clamped neurons.

### Multi-electrode Arrays (MEAs)

MEAs recordings were performed in two independent laboratories with slightly different equipment and protocols. The two methods differed as follow:

### Method #1 used for MEA experiments (Figures 4, 6 performed in Bardy Lab)

***Cell culture on MEA plates #1:*** The neuronal cultures were generated from NPCs in the neural maturation medium (NMM) as described above. Fourteen days after neuronal maturation in NMM, cultures were dissociated using Accutase and resuspended in NMM at 15,000 viable neurons/μL. A 10 μL droplet of cell suspension was added directly over the recording electrodes of each well of the MEA, which were previously coated with 10 μg/mL poly-L-ornithine (Sigma) and 5 μg/mL laminin (Thermo Fisher Scientific). Sterile water was added to the area surrounding the wells, and the MEA was incubated at 37°C, 5% CO_2_ in a cell culture incubator. After 1 hour, 300 μL NMM was gently added to each well. Half of the culture medium was exchanged with fresh medium three times per week. Fifty-two days after maturation commenced, the medium was replaced with one of three different media: BrainPhys™ Imaging, ACSF or BrainPhys™ (STEMCELL Technologies), each supplemented with the midbrain or cortical basal media supplements, as described above. ***MEA recording and analysis #1:*** MEA recordings were taken by a Maestro Pro MEA system (Axion Biosystems). Lumos MEA 48 plates (Axion Biosystems) were maintained at 37°C with 5% CO_2_ environment during recordings. Recordings were not made for at least 10 minutes after transferring a plate into the MEA. Version 2.4 AxIS acquisition software (Axion Biosystems) was used to sample voltage potentials simultaneously across 16 electrodes per well. The sampling frequency was 12.5 kHz. The threshold for detecting spikes was set on a per-electrode basis and was defined as the voltage exceeding 6 standard deviations away from the mean background noise‥spk files were analyzed using NeuralMetricTool version 2.5.1 software (Axion Biosystems) and custom MATLAB scripts (MathWorks). An electrode was defined as active with a mean firing rate greater than 0.017 Hz (> 1 spike per min). ***Optogenetics on MEA #1:*** Optical stimulation parameters were configured using Stimulation Studio software in AxIS. Optical stimulation was performed by a Lumos optical stimulator (Axion Biosystems). The Lumos stimulation protocol was as follows: Every 20 seconds, wells were simultaneously exposed to a stimulation burst consisting of 10 flashes of blue (475 nm) light, each lasting 10 ms with a 100ms interval between each flash. Lumos blue LED stimulation at 100% intensity is >3.9mW/mm^2^ according to the manufacturers. The intensities of stimulation used here ranged from 5-50%.

### Method #2 used for MEA experiments (Figure 7, performed in SCT Lab)

***Cell culture on MEA plates #2:*** Frozen neural progenitor cells (NPCs) induced pluripotent (iPS) stem cells (previously characterized in ^51^) were differentiated into neuronal precursors with the STEMdiff™ Neuron Differentiation Kit (STEMCELL Technologies). Neuronal precursor cells were then frozen with Cryostor10 (STEMCELL Technologies) in liquid nitrogen storage and thawed later for experiments. Post-thaw, cells were plated at 10,000 cells in a 40μL droplet of STEMdiff™ Neuron Differentiation medium in a Cytoview MEA 48 plate (M768-tMEA-48B, Axion Biosystems) pre-coated with 15μg/mL poly-L-ornithine (Sigma) and 10μg/mL laminin (Sigma) (12 wells total). After a 2-hour incubation in a temperature-controlled incubator, 150μL of the medium was added to each well. On post-thaw day 1, 150μL of complete BP was added to each well. One well of the total 12 was eliminated from testing due to poor cell adhesion. Cells were cultured for 6 weeks in complete BP with half-media changes performed every 3-4 days. Subsequently, media in 5 out of 11 of the wells was exchanged with complete BPI for a period of 2 weeks before switching back to BP. ***MEA recording and analysis #2:*** Spontaneous neuronal activity was acquired at 37°C under a 5% CO_2_ atmosphere using a Maestro system (Axion Biosystems) at a sampling rate of 12.5 kHz/channel. For recording, a Butterworth band-pass filter (200 Hz - 3000 Hz) was applied and the adaptive threshold spike detector was set at 6X standard deviation. A 15-minute recording was taken twice a week. Data from the last 10 minutes of each recording was exported for analysis using AxIS software (2.5.1; Axion Biosystems). An electrode was defined as active with a mean firing rate greater than 0.017 Hz.

### Statistics

Statistical analyses were performed with Graphpad Prism 8. Statistical significance was assessed between different media and BrainPhys™ Imaging groups using the following: two-tailed non-parametric unpaired (Mann Whitney) tests, unpaired t-tests, Wilcoxon signed-rank test for non-parametric paired data sets, and sum-of-squares F-test.

### Ethics

All patient tissues collected, and experimental procedures were performed in accordance with The Women's and Children's Health Network HREC/17/WCHN/70. Human cells were collected from patients with informed consent. Human embryonic stem cells (WA09, H9) were obtained from WiCell (agreement 20-W0500).

## Data availability statement

The data that support the findings of this study are available from the corresponding author upon reasonable request.

## Author Contributions

CB, CM, VL, and EK designed the project. KM, BM, RA and MZ executed and analyzed the fluorescent phototoxicity experiments. MZ, KM, AA, LC, AH and AM performed the absorbance and fluorescence assays. BM, RA, CM, MZ and KM performed the MEA experiments. KM, CM, BM, JT and MvdH performed tissue culture. MZ, and AS executed and analyzed the patch-clamping, calcium imaging, and optogenetics experiments. AMJ and PR performed the absorbance analysis. CB wrote the manuscript with contributions from MZ and KM and feedback from all authors.

## Acknowledgments

We thank Sebastian Loskarn and Zarina Greenberg for technical assistance. This work was generously supported by Perpetual Impact Philanthropy, the Brain Foundation, Rebecca L. Cooper Foundation, Ian Potter Foundation, Boileau Corporate Philanthropy, Parkinson’s South Australia, MRFF Australian Government and Department of Health, Australian Research Council LIEF grant (to CB); Centre for Nanoscale and Biophotonics (CE140100003), Future fellowship (FT180100565) (to MRH), the US National Research Council Senior Fellow Program to the US Air Force Research Laboratory (to AMJ), the Netherlands Organisation for Scientific Research Rubicon Fellowship 019.163LW.032 (to MvdH), and the RMIT University Vice-Chancellor’s Research Fellowship (to PR) and The Australian Government Research Training Program Scholarship (to MZ, BM, RA). Flow cytometry analysis and cell sorting were performed at the South Australian Health Medical Research Institute (SAHMRI) in the ACRF Cellular Imaging and Cytometry Core Facility. The Facility is generously supported by the Detmold Hoopman Group, Australian Cancer Research Foundation and Australian Government through the Zero Childhood Cancer Program.

