## Supplementary figures for "BrainPhys neuronal medium optimized for imaging and optogenetics in vitro"

**Figure S1 (related to Figure 1 and 2): Influence of BrainPhys Imaging on autofluorescence and signal-to-background ratio compared to competitive media.**

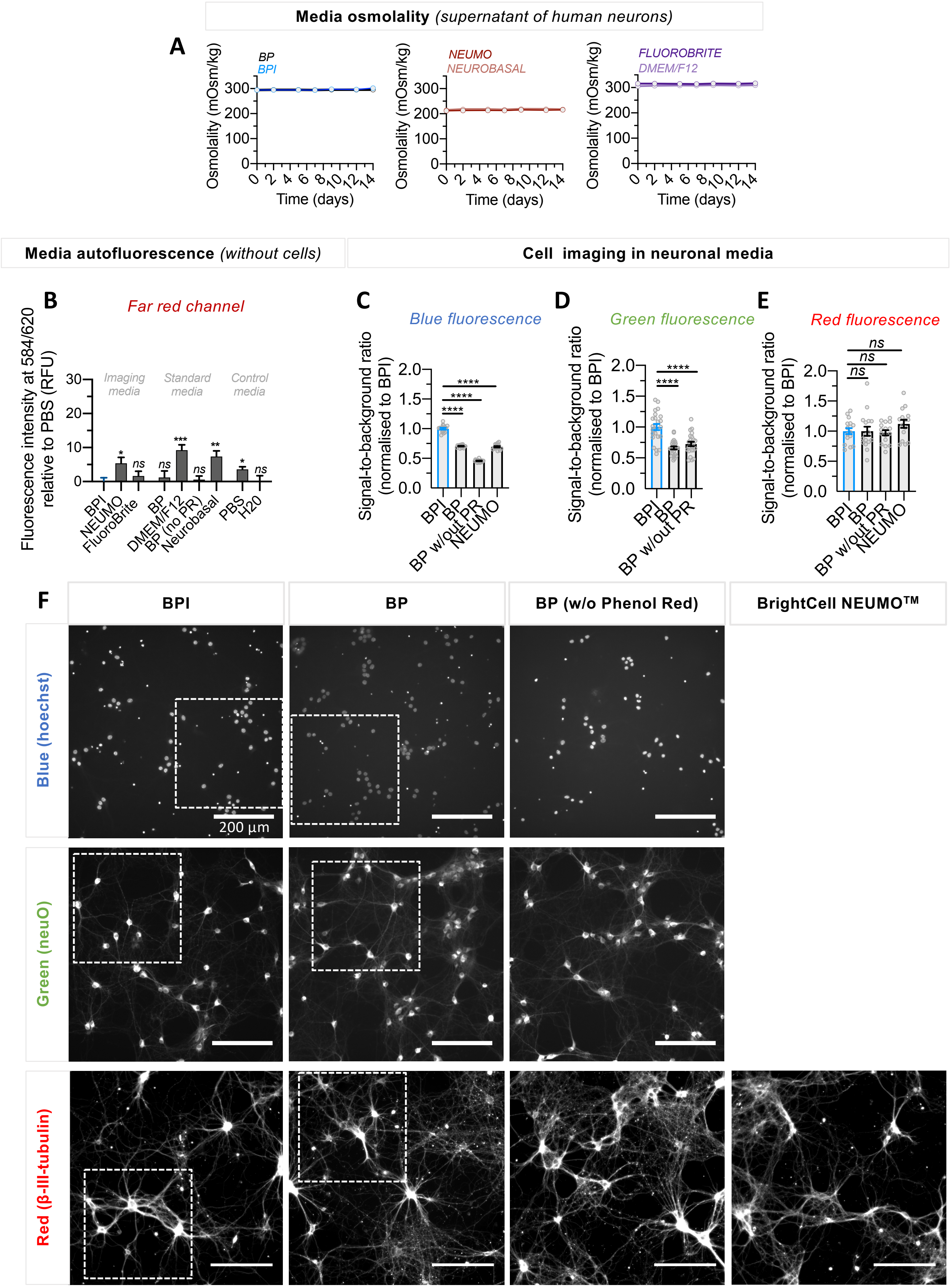

**A,** shows the osmolality of supernatant collected from human cortical iPSC-derived neurons matured in BP > 12 weeks and then switched to either BPI, BP, NEUMO, Neurobasal, DMEM/F12 or FluoroBrite basal media for 14 days. The same supplements for cortical neurons were added to all conditions. BPI maintained osmolality measurements over the 14 days at physiological levels. For each time point, results were generated from four replicate wells per medium. **B**, far-red autofluorescence intensity of imaging, standard and control media collected in a 96-well plate reader set to an excitation/emission of 584/620 nm. Results were generated from eight replicate wells per medium. For normalization, the mean fluorescence intensity in PBS was subtracted from the other media. **C-E**, the signal-to-background ratio quantification of live rat cortical primary neurons labelled with NeuroFluor NeuO and fixed neurons stained with β-III tubulin or Hoechst 33342 and imaged in BPI and BrainPhys (standard or without phenol red) and NEUMO. For BPI, the signal-to-background ratio of Hoechst 33342 **(C)** was significantly higher compared to BP, BP (without Phenol Red) and NEUMO. Similarly, BPI also improved image quality relative to BP and BP (without Phenol Red) for neurons labelled with NeuO **(D)**. No significant differences were found for β-III tubulin between test media **(E)**. Across Hoechst 33342, NeuO and β-III tubulin channels; a total of 16, 25 and 16 respective field-of-views were analyzed from one well per condition. Data was collected from two biologically independent experiments and shown normalised to BPI. For further image quantification and parameters details, refer to 'methods'. **F,** representative full-scale images of primary rat cortical neurons displayed in **figure 2E**. White boxes represent cropped images shown in Figure 2E. Images captured using a 20x air immersion lens (0.45 NA) are displayed with the following maximum/minimum intensity counts across all test media: 0/8500 (Hoechst), 0/3600 (NeuO), 1000/12000 (β-III tubulin). Values in **(B-E)** represent mean±SEM, significance determined via two-tailed non-parametric unpaired (Mann Whitney) tests. P-values are annotated as follows: **** for $P<0.0001$ and ns for $P>0.05$.

**Figure S2 (related to figure 2). Influence of BrainPhys Imaging on the signal-to-background ratio of live human neurons.**

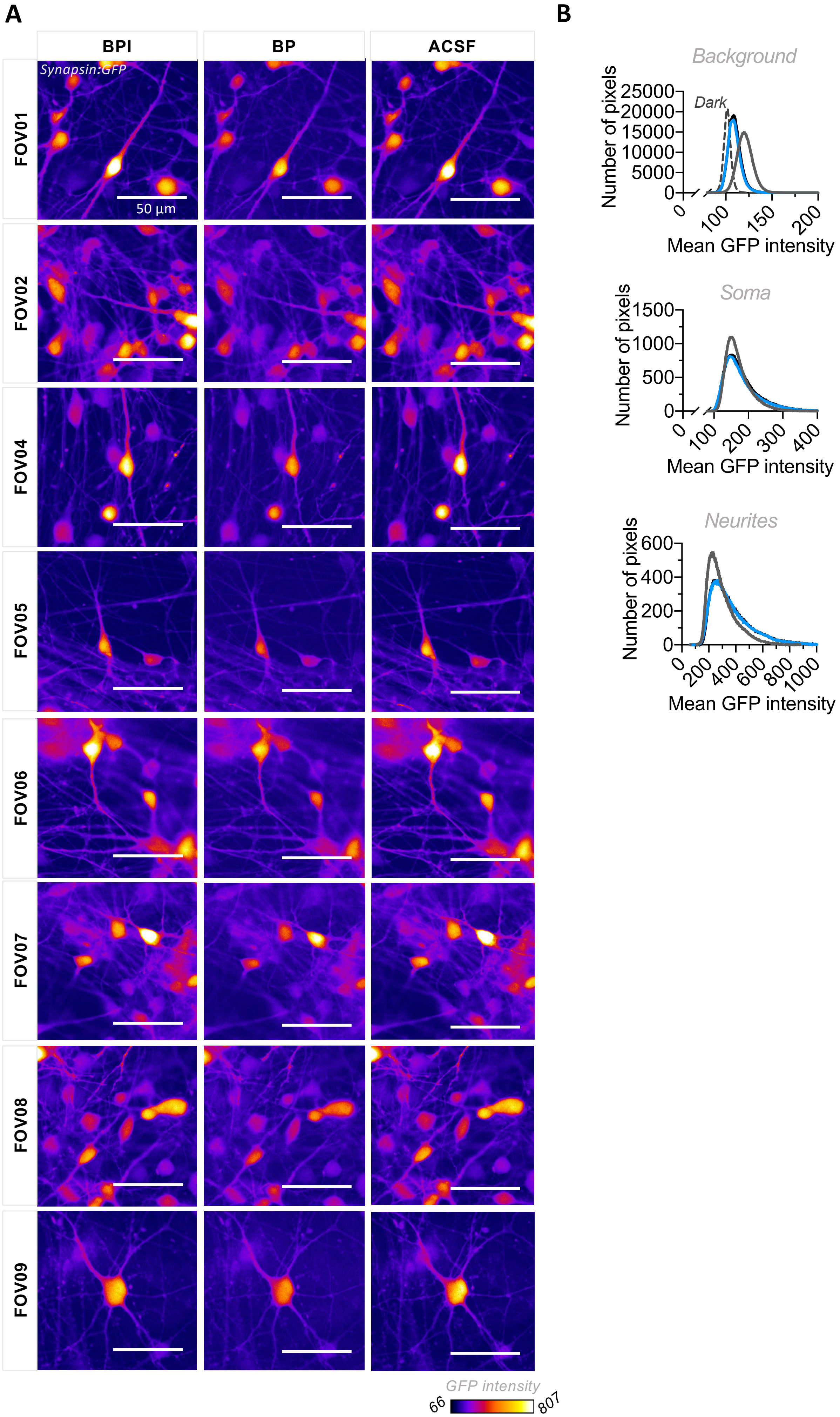

**A,** representative fluorescent images of live human iPSC-derived neurons expressing GFP (excitation/emission peaks: 488/509 nm; filters: 470±40/525±50 nm) and imaged in BPI, BP or artificial cerebrospinal fluid (ACSF) using a 40x water immersion lens (0.8 NA). Field-of-views (FOV) 1,2,4-9 are shown, and were analysed for figure **2B-C and S2B.** Higher mean GFP intensities at soma (yellow) and neurite (magenta) regions can be witnessed in BPI and ACSF relative to BP, whereas background intensities were reduced in BPI and ACSF (purple). **B,** shows the number of pixels at background, soma and neurite regions imaged in BP (grey), BPI (blue), and ACSF (black) across a mean GFP intensity range. Camera dark counts shown as 'dark' (top panel). Analysis was conducted on GFP images from 9 field-of-views of the same coverslip.

**Figure S3 (related to figure 2). Representative images of live human neurons in BrainPhys Imaging compared to ACSF and BrainPhys.**

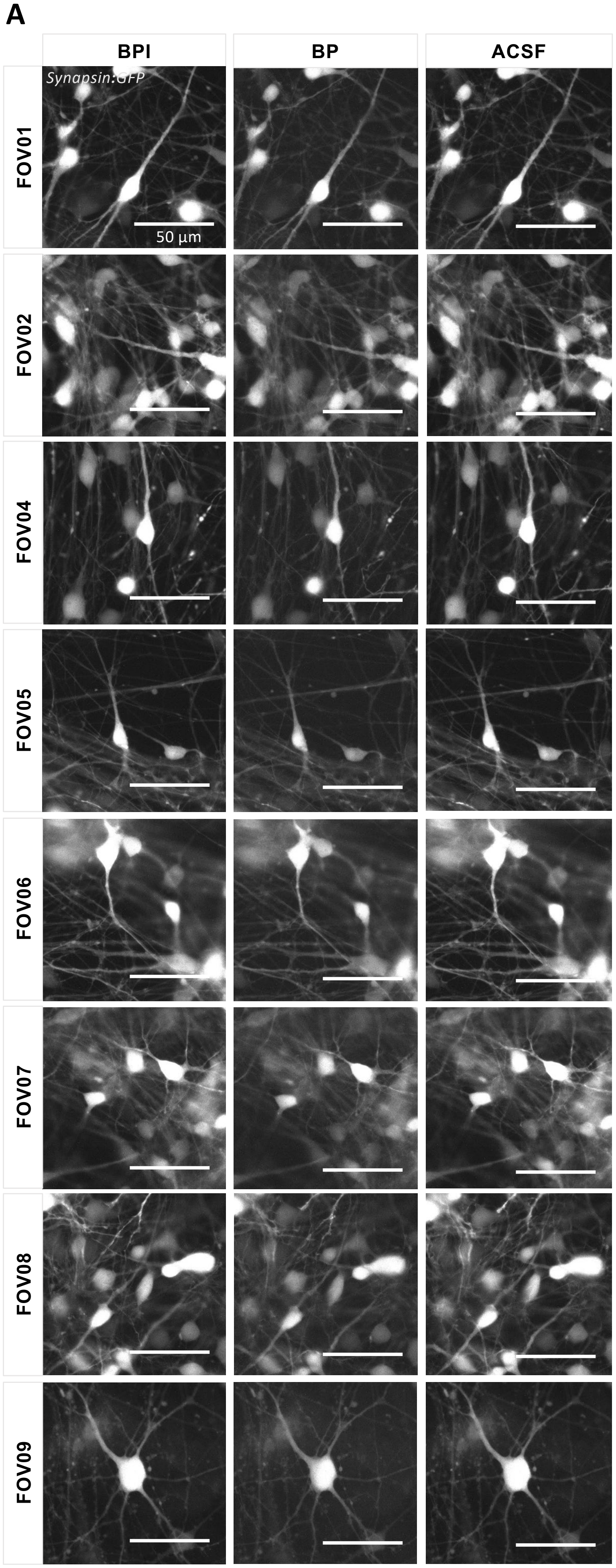

**A,** representative gray-scale images of live human iPSC-derived neurons expressing GFP (excitation/emission peaks: 488/509 nm; filters: 470±40/525±50 nm) and imaged in BPI, BP or artificial cerebrospinal fluid (ACSF) using a 40x water immersion lens (0.8 NA). Field-of-views (FOV) 1,2,4-9 are shown, and were analysed for figure **2B-C and S2B.** Images are displayed with maximum/minimum intensity counts of 75/400 across all test media.

### Figure S4 (related to figure 3). BrainPhys Imaging maintains cell viability following light exposure.

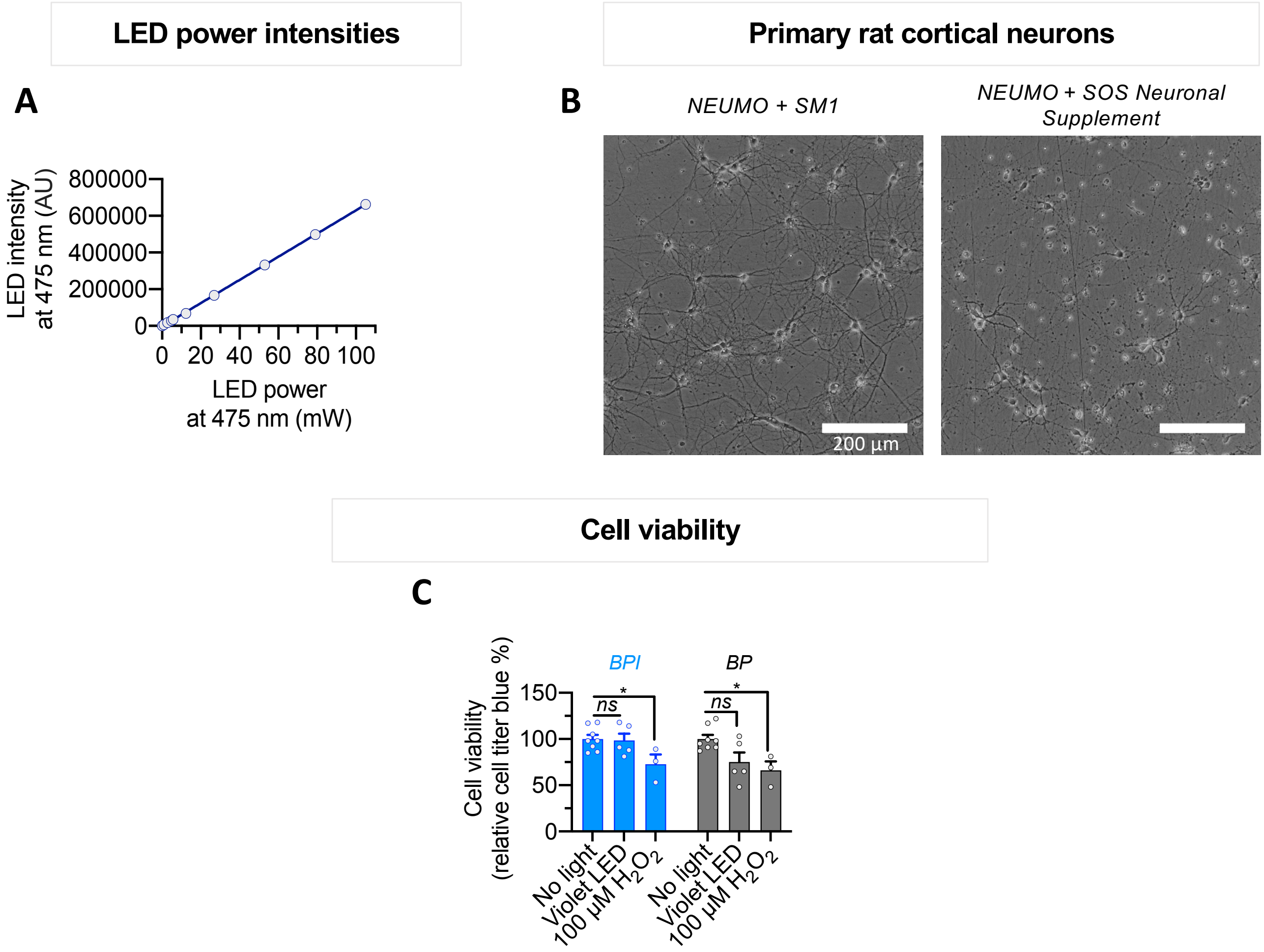


**A,** LED intensity recorded from a single LED on a LUMOS optical simulator (at 475 nm), and its corresponding LED power (mW). **B**, primary rat cortical neurons could not be sustained for 11 DIV in NEUMO supplemented with SOS compared to NEUMO supplemented with SM1. **C,** the cell viability (CellTiter-Blue® reagent) of primary rat cortical neurons was assessed following violet LED exposure inside a temperature-controlled incubator for 1 hour and treatment with 100 μM of $H_2O_2$. Data was collected across 3-5 independent experiments across 3 biological replicates. Results are shown normalised to 'no light' or 'control' conditions. Cell viability in BPI or BP media supplemented with SM1 following violet LED light exposure was maintained across all conditions but significantly reduced after $H_2O_2$ treatment in BPI and BP media. Values are shown as mean ± SEM. Significance determined via two-tailed non-parametric unpaired (Mann Whitney) tests **(C)**. P-values are annotated as follows: * for $P<0.05$, and ns for $P>0.05$.

**Figure S5 (related to figure 3): BrainPhys Imaging reduces short term phototoxicity of neuronal cultures.**

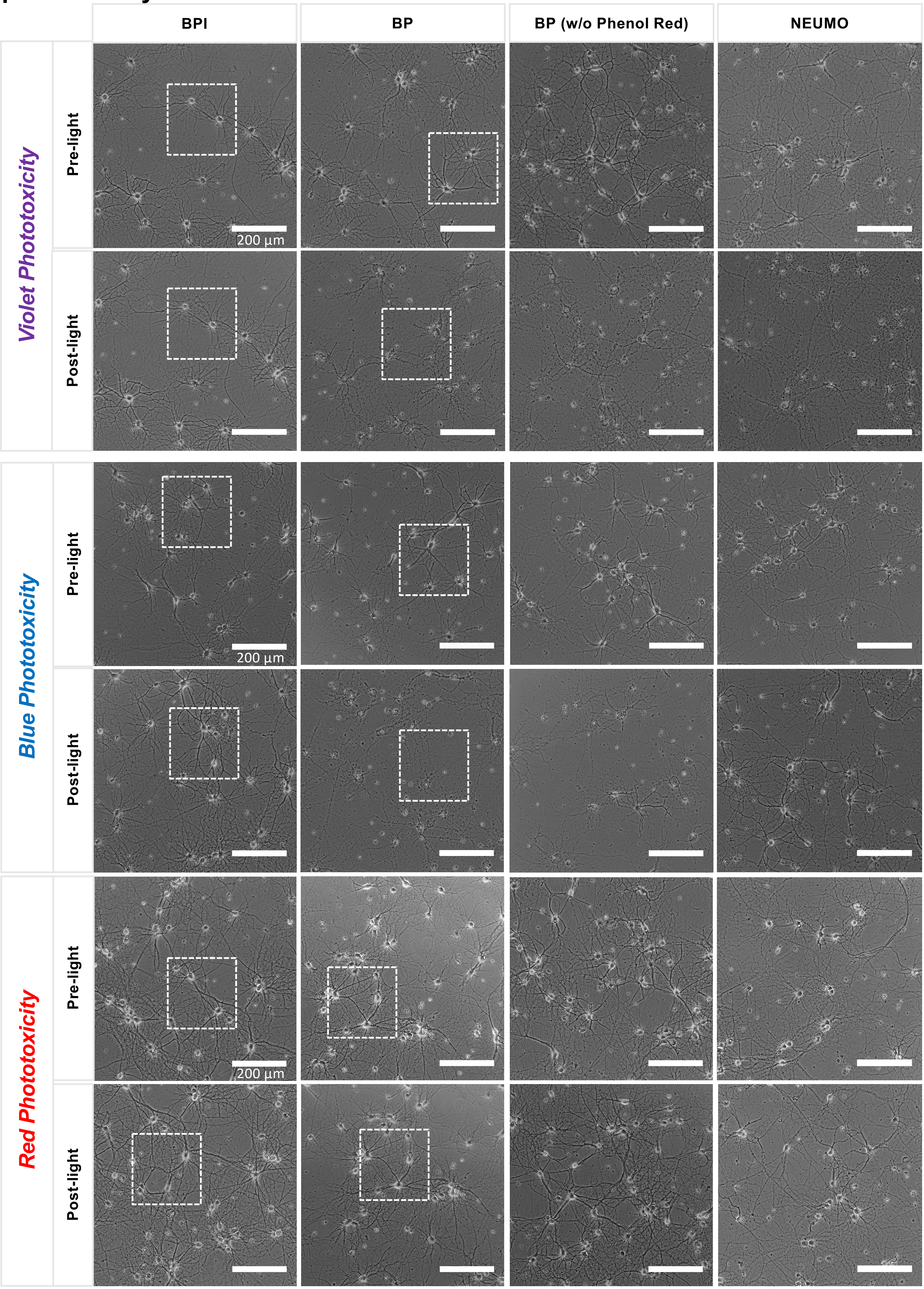

Representative images of rat cortical primary neurons cultured in standard BrainPhys, BPI, BrainPhys without phenol red, and NEUMO before and after exposure to red, blue and violet LED light. White boxes represent cropped images shown in figure 3A-C.

**Figure S6. (related to figure 4) Relative to ACSF, BrainPhys Imaging establishes equivalent electrophysiological activity in human neurons.**

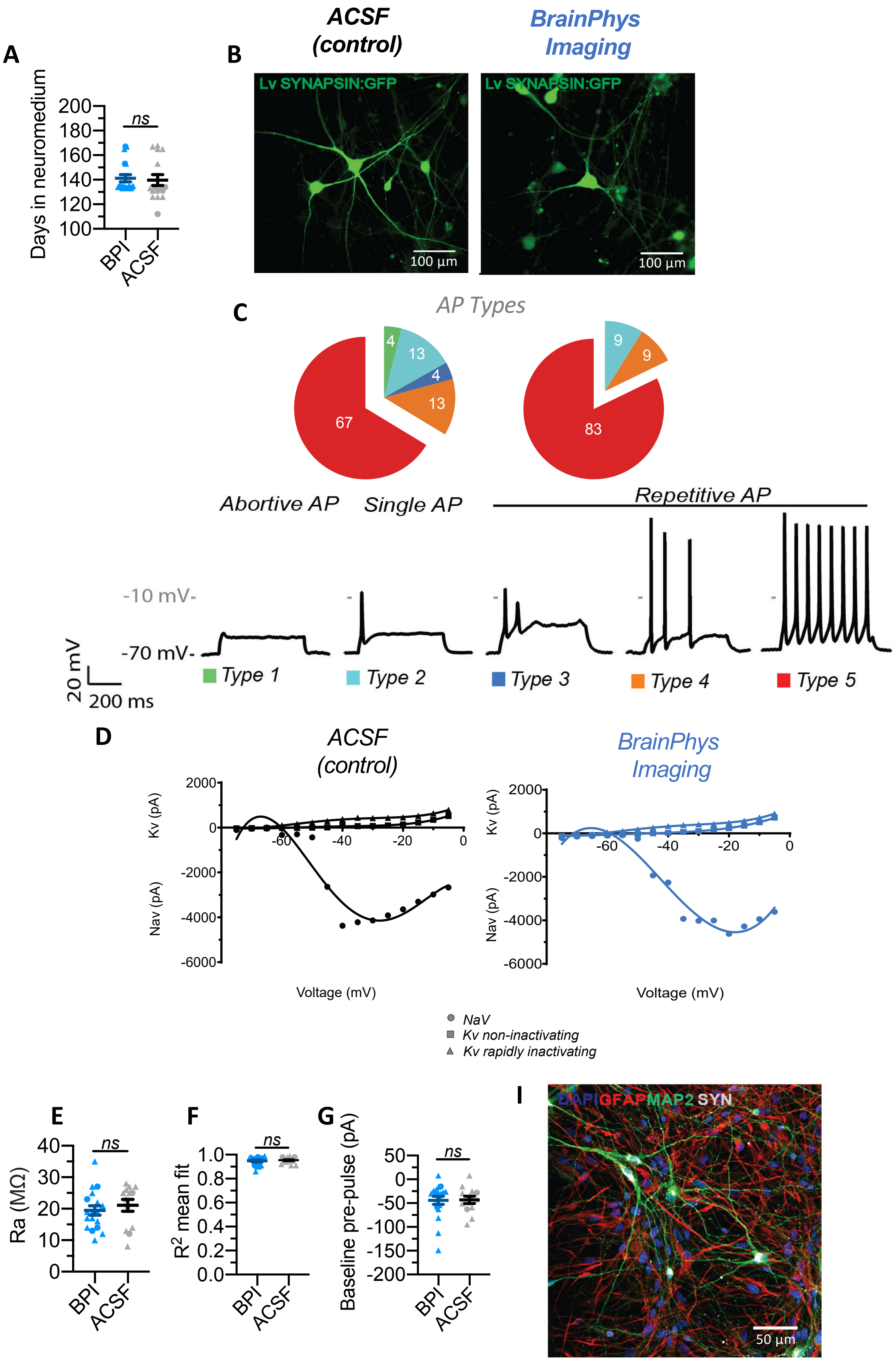

**A,** the maturation time (days) of type 5 human neurons analysed and recorded across eighteen coverslips, incubated in BrainPhys + supplements ≥ 112 days (37degC, 5% $CO_2$ and 21% $O_2$) and then patched in BPI (n=19 neurons) or ACSF (n=16 neurons). **B,** shows images of neurons in ACSF or BPI expressing synapsin (GFP) before patch-clamping. **C,** the proportion of action potentials (APs) types evoked in human neurons when patched in BPI or ACSF with corresponding exemplary traces. Data collected from a total of 47 neurons across eighteen coverslips in BPI (n=23) or ACSF (n=24). Type 5 neurons (evoked APs >10 Hz; >-10 mV peak amplitudes), type 4 neurons (evoked APs <10 Hz, >-10mV peak amplitude), type 3 neurons (>1 evoked AP and >1 aborted spikes < -10mV peak amplitudes), type 2 neurons (>1 evoked AP >-10mV amplitude, followed by a plateau), type 1 neurons (not able to fire APs > -10mV). **D,** analyzed current-voltage (IV) curves of selected type 5 human neurons patched in ACSF or BPI medium, showing the relationship between voltage-dependent Na+ and K+ (non-inactivating and rapidly inactivating) currents (pA) and depolarizing current steps (mV). Voltage clamped at -70 mV with +5 mV steps. **E-G,** quantification of access resistance (Ra), baseline pre-pulse and $R_2$ mean fit, for all type 5 neurons. Symbols represent human neurons tested first (triangles) or second (circle) in either medium (**A, E-G**)**. I,** shows an example of immunostaining of human neuronal cultures matured in BrainPhys + supplements > 4 weeks. DAPI (blue), GFAP (red), MAP2 (green), Synapsin (grey) highlighting neuronal and astrocyte populations along with dendritic projections. Values are shown as mean ± SEM. Significance determined via two-tailed non-parametric unpaired (Mann Whitney) tests **(A,E-G)**. P-values are annotated as follows: ns for P>0.05.

**Figure S7. (related to figure 4) Relative to BrainPhys, BrainPhys Imaging supports equivalent electrophysiological activity in human neurons.**

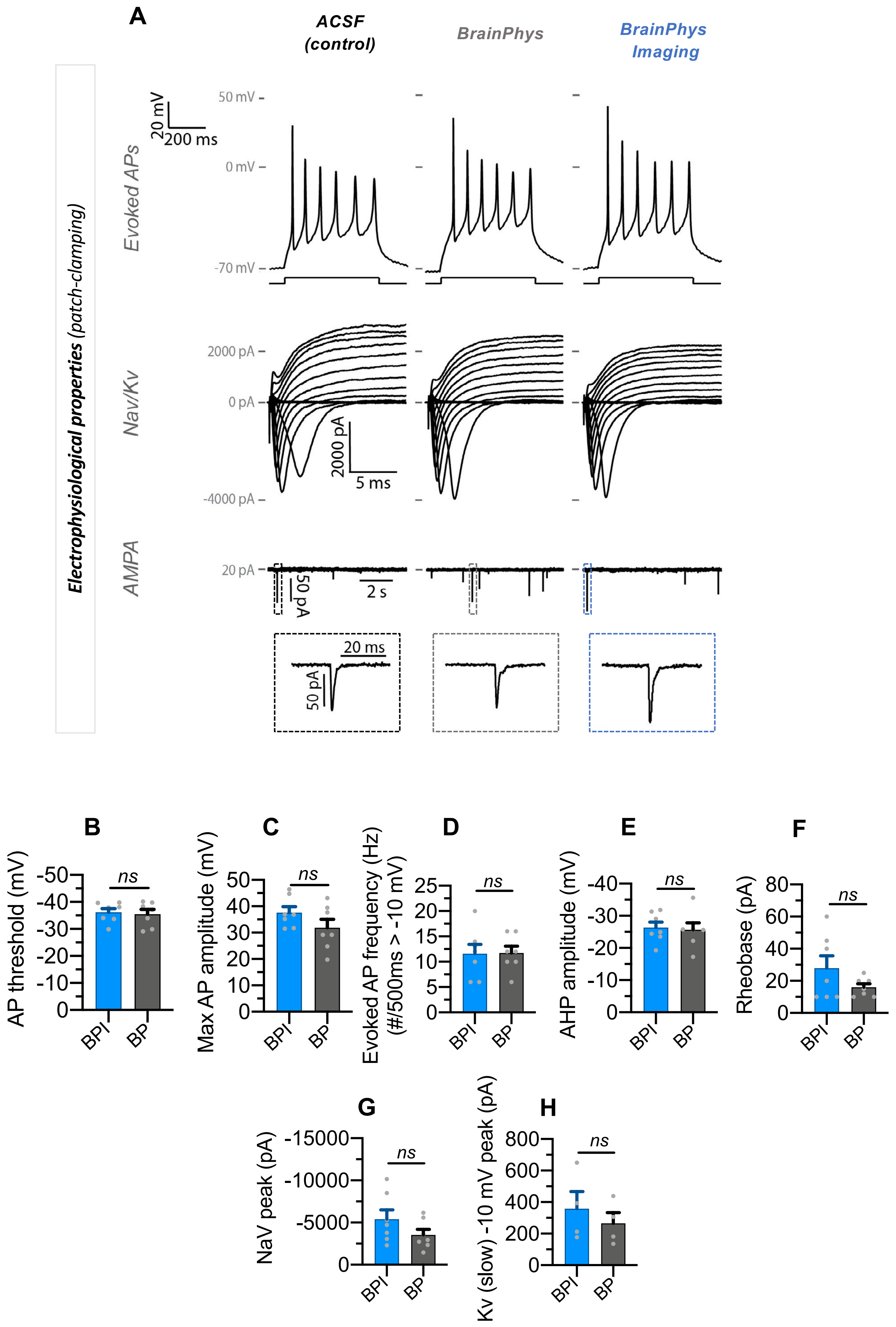

Single-cell patch-clamp recordings of human pluripotent stem cell-derived neurons matured in standard BP + supplements medium for >12 weeks, and patch clamped in BP or BPI media. All patch clamped cells (n=8) included in the analysis were classified as Type-5 neurons (evoked APs > 10 Hz, amplitudes > -10 mV) based on methods previously established in Bardy et al. 2016. See also **Figure 4** and **S6**. A subset of neurons (n=7) was recorded in both media. All neurons were patched from a total of 4 coverslips. Each point on the graphs in panels B-H represents a single neuron. **A, top:** shows similar evoked action potential (AP) traces following a 500 ms depolarizing current step for the same neuron patched in ACSF, BrainPhys and BPI. **Middle:** corresponding current-voltage characteristics (I-V curve) reveal similar voltage-dependent sodium (Nav) and potassium (Kv) current amplitudes for the same neuron patched across ACSF, BrainPhys or BPI. Nav and Kv current traces shown, respectively, below and above the x-axes. Current steps of +5 mV increments were used from resting potential at -70 mV. **Bottom:** shows typical spontaneous ePSC traces from the same neuron patch clamped in ACSF, BrainPhys and BPI. No significant differences were found between the electrophysiological properties of human neurons patch-clamped in either BP (n=7) or BPI (n=7) basal media for **(B)** AP thresholds, **(C)** peak AP amplitudes, **(D)** firing frequencies of AP evoked by 500ms depolarization steps (spikes with amplitudes > -10 mV included), **(E)** peak afterhyperpolarization (AHP) amplitudes, **(F)** rheobase values, and (**G)** peak NaV current amplitudes. **H,** peak amplitudes of slowly inactivating Kv currents in BPI (n=4) and BrainPhys (n=4) were also similar. Values are shown as mean ± SEM. Significance determined via two-tailed non-parametric unpaired (Mann Whitney) tests **(B-G)** and non-parametric paired (Wilcoxon) tests **(H)**. P-values are annotated as follows: ns for P>0.05.

**Figure S8: (related to figure 5) Calcium imaging of human neurons in BrainPhys Imaging versus ACSF shows similar proportions of wave and spike events.**

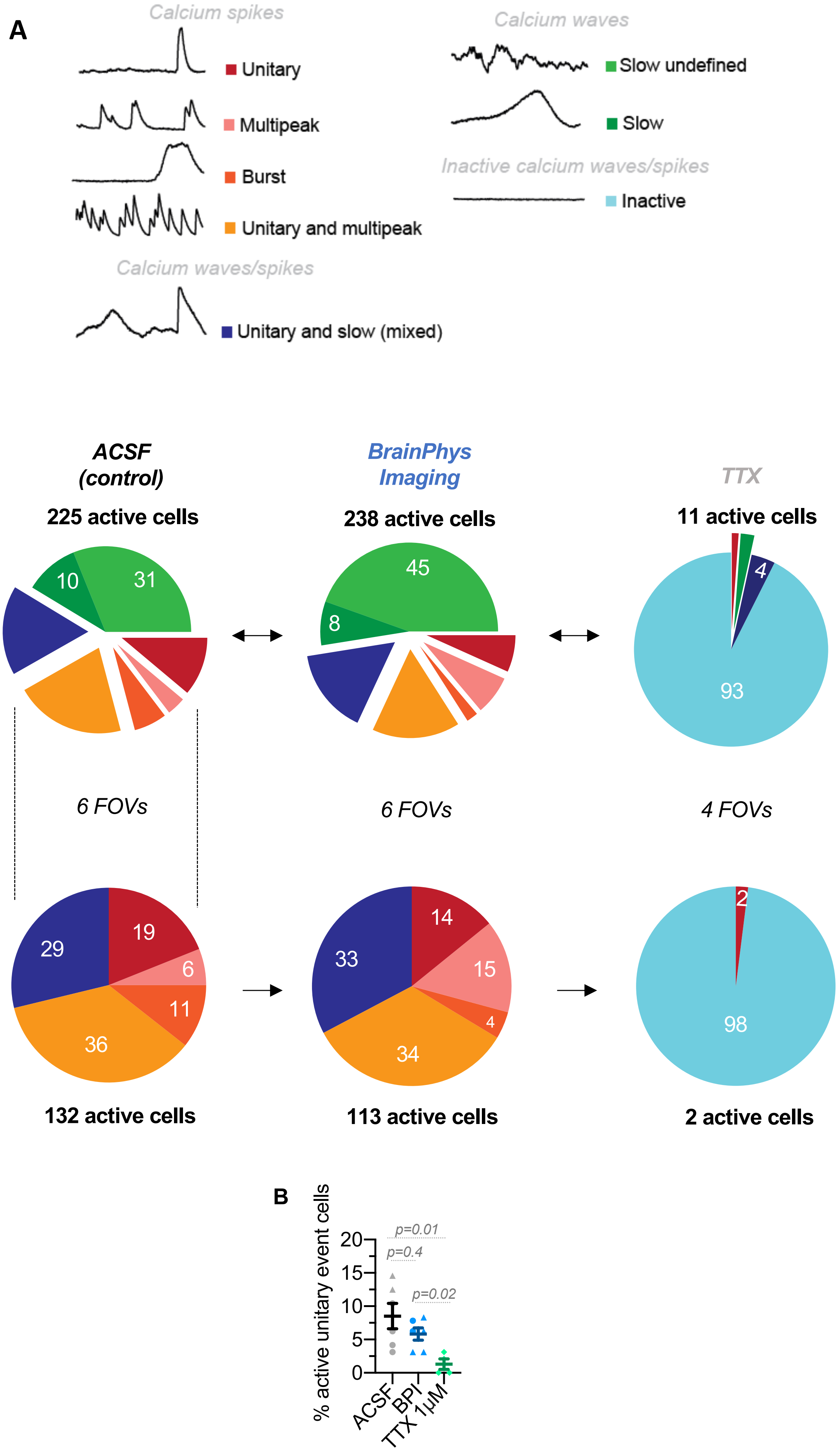

**A,** shows representative calcium image-sequence traces filmed at 5Hz from human neurons in BPI or ACSF. Events were categorized into calcium spikes (fast-rising phase events), calcium waves (slow rising phase events), calcium waves/spikes (a combination of fast and slow rising phase events) or inactive events. *Top pie chart:* calcium imaging data collected across six field-of-views (FOV) from two coverslips in ACSF (n=225 cells), BPI (n=238 cells) and TTX (n=11 cells) were categorized into calcium event types. *Bottom pie chart:* a similar distribution of calcium spike event types was witnessed in selected cells initially displaying active calcium spikes in ACSF (n=132 cells) then switched to BPI (n=113 cells) and TTX (n=2 cells) perfusates. TTX perfusion showed a significant reduction in active spike events. Note that cells inactive across all three perfusates were excluded from the analysis. **B,** quantification of the mean percentage of cells with unitary events witnessed per FOV in different perfusates. Symbols in **B** represent the order of media perfusion for each FOV: first (triangle), second (circle), or last (rotated square). **B,** data is mean±SEM. Significance determined via two-tailed non-parametric unpaired (Mann Whitney) tests.

**Figure S9: (related to figure 5) Influence of BrainPhys Imaging medium on the activity of human neurons compared to BrightCell™ NEUMO and FluoroBrite™ DMEM.**

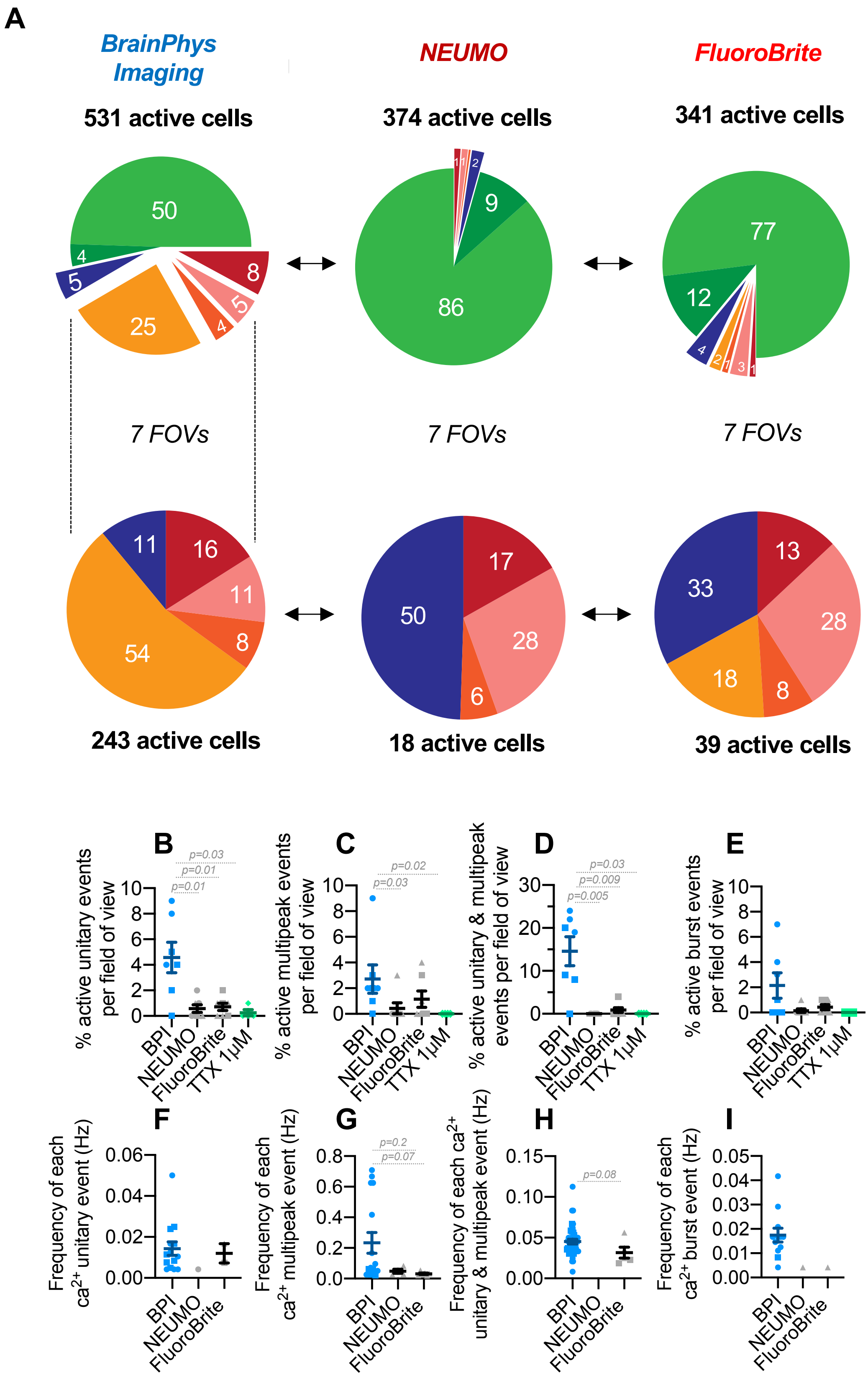

**A**, *Top pie-chart:* breakdown of time-lapse image sequences (filmed at 5Hz, 1200 frames) measuring intracellular calcium events (Fluo-4 calcium sensor) categorized into active calcium spikes, waves, and spikes/waves. A total of 909 cells across 7 field-of-views (FOV) were recorded, and inactive cells across all three perfusates were excluded from further analysis. Remaining active cells in BPI (n=531 cells), NEUMO (n=374 cells) or FluoroBrite (n=341 cells) perfusates across two coverslips were selected for and analysed. *Bottom pie chart:* percentage breakdown of active calcium spikes and spike/wave events when imaged in either BPI (n=243 cells), NEUMO (n=18 cells) and FluoroBrite (n=39 cells) perfusates. **B-I**, Quantification of the mean percentage (B-E) and frequency (F-I) of active unitary, multipeak, unitary and multipeak, and burst events per FOV in the different media. Note that the addition of TTX (1 µM) perfusate completely blocked all unitary spike event activity. Symbols in **B-I** represent the order of media perfusion for each FOV: first (triangle), second (circle), third (square) or last (rotated square). **B-I,** data is mean±SEM. Significance determined via two-tailed non-parametric unpaired (Mann Whitney) tests.

**Figure S10: (related to figure 6) BrightCell™ NEUMO limits the basic firing properties of human neurons compared to BrainPhys Imaging.**

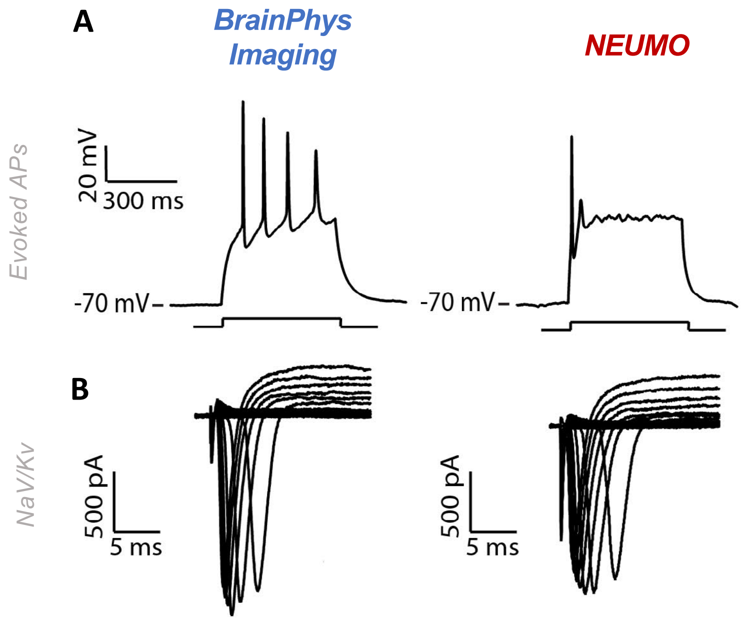


Example single-cell recording from a type 4 human neuron patch-clamped in BPI or BrightCell™ NEUMO perfusate. **A**, shows typical evoked action potential (AP) traces following a 500ms depolarising current step when patched and held at -70mV in either perfusate. **B**, Current-voltage characteristics (I-V curve) of a human neuron patched in either perfusate when held at −70 mV with +5 mV current steps. Voltage-dependent sodium (NaV) and potassium (KV) current traces shown, respectively, below and above the x-axes.
